# Vaccination by single dose sporozoite injection of blood stage attenuated malaria parasites

**DOI:** 10.1101/2023.10.24.563045

**Authors:** Julia M. Sattler, Lukas Keiber, Aiman Abdelrahim, Xinyu Zheng, Martin Jäcklin, Luisa Zechel, Catherine A. Moreau, Manuel Fischer, Chris J. Janse, Angelika Hoffmann, Franziska Hentzschel, Friedrich Frischknecht

## Abstract

An efficient malaria vaccine remains elusive. As an alternative to malaria subunit vaccines, vaccination approaches are currently explored using live *Plasmodium* parasites, either attenuated mosquito-derived sporozoites or attenuated blood stage parasites. Both approaches would profit from the availability of attenuated and avirulent parasites with a reduced blood stage multiplication rate. Ideally, such slow growing parasites would proceed normally through the mosquito but cause a self-limiting infection upon transmission. Here we screened gene-deletion mutants of the rodent parasite *P. berghei* and the human parasite *P. falciparum* for slow growth. In addition, we tested the *P. berghei* mutants for avirulence in mice and self-resolving blood stage infections, while preserving sporozoite formation and liver infection. Targeting fifty genes yielded seventeen *P. berghei* gene-deletion mutants with two mutants causing self-clearing infections in mice while retaining full transmissibility through mosquitoes. For those, infection of mice by a low number of blood stages, infected-mosquito bites or by single injection of sporozoites led to protection from disease after challenge with wild type sporozoites. Two of six generated *P. falciparum* gene-deletion mutants showed a slow growth rate. Slow growing, avirulent *P. falciparum* mutants will constitute valuable tools to inform on the induction of immune responses and aid in developing new as well as safeguarding existing attenuated parasite vaccines.

## Introduction

Malaria is caused by the cyclical infection of red blood cells by *Plasmodium* parasites, which undergo a complex life cycle (Figure 1a) and are first injected into the vertebrate by a mosquito bite. During the bite *Plasmodium* sporozoites are deposited in the skin where they rapidly migrate to enter blood vessels to be transported to the mammalian liver where they invade hepatocytes and develop into red blood cell (RBC) invading merozoites. Different parasite life cycle stages, such as sporozoites, liver and blood stages, can induce immune responses, which have been exploited for the generation of subunit or whole parasite vaccines ^1–3^. The most advanced subunit malaria vaccine developed to date, RTS,S (Mosquirix), is currently in phase 4 clinical development and offers 30-50% protection against clinical malaria ^4,5^ with the newer R21 recently showing around 75% protection ^6^. These so-called subunit vaccines are composed of part of the malaria parasite circumsporozoite protein, the most abundant protein on the outer surface of the sporozoite ^7,8^. Several other subunit vaccines are also being developed and evaluated ^9–11^.

**Fig. 1.**
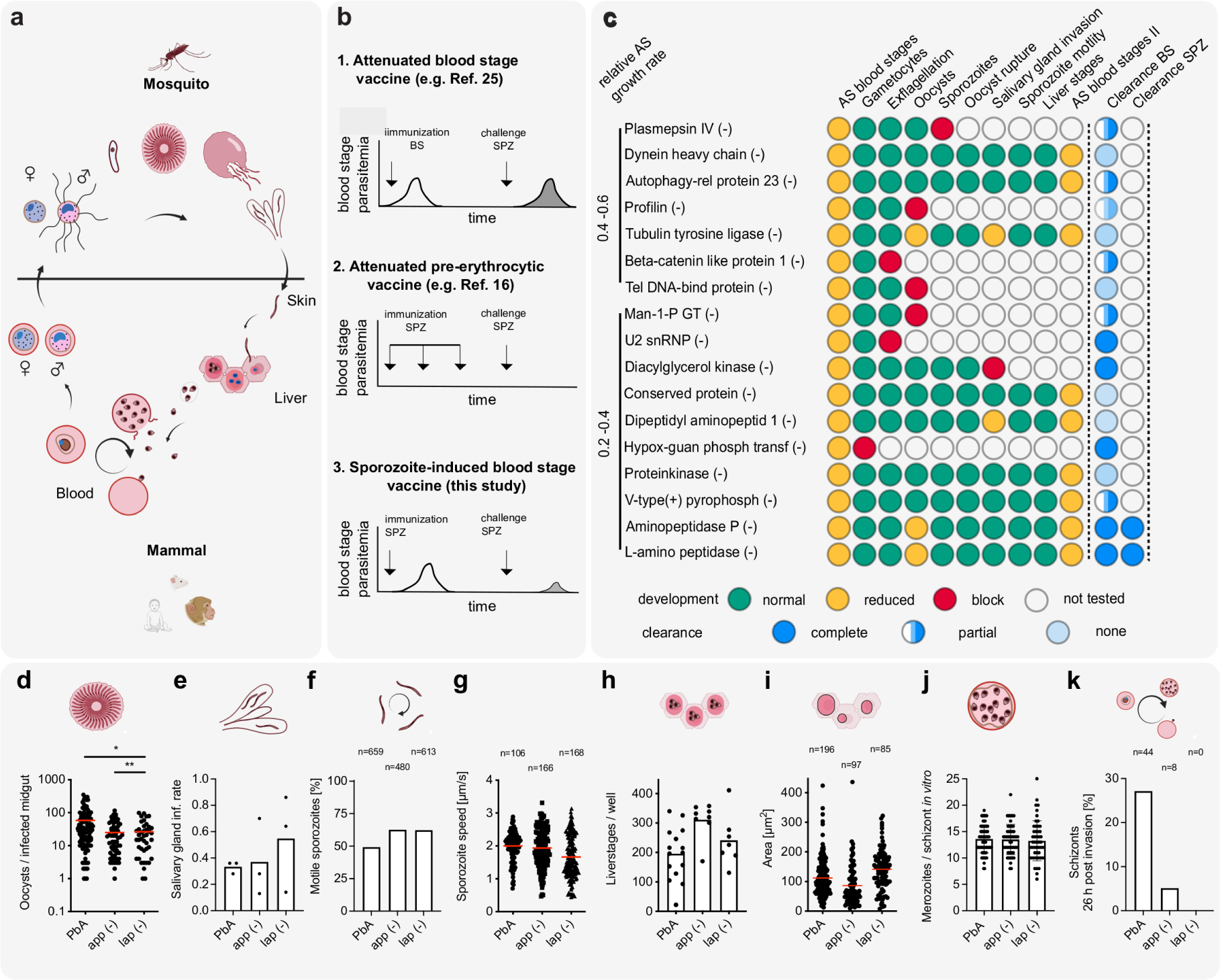
Screening for *P. berghei* gene-deletion mutants with slow growing, blood stages that retain mosquito transmissibility. **a)** *Plasmodium* life cycle indicating the stages investigated in this report. **b)** Schematics of genetic attenuation strategies. Mutant parasites can be generated to (i) attenuate in the blood stage, (ii) enter into the liver to arrest their development and, as shown here (iii) complete the life cycle but attenuate in the blood stage. Immunization schemes and model blood stage parasitemia curves upon immunization and challenge are shown. **c)** Life cycle progression analysis of gene-deletion mutant lines with reduced blood stage growth lacking the indicated genes. Top to bottom: fastest to slowest blood stage growth rate of the parasite lines. Green and yellow circles: similar and reduced developmental characteristics, respectively compared to wild type. Red circles: a complete block in life cycle progression. Blue circles: relative level of clearance of blood stages after infection by 100 iRBC *i.v.* or by sporozoite-induced infections (1.000 spz *i.v.* or natural transmission) in C57Bl/6 mice. AS: asexual, BS: blood stage, SPZ: sporozoites. **d)** Oocyst numbers in mosquitoes infected with wild type (*Pb*A), *app(-)* and *lap(-)*. * and ** indicate p<0.05 and p<0.01 respectively; a Shapiro-Wilk normality test was applied followed by a Kruskall-Wallis test. **e)** Salivary gland invasion in mosquitoes infected with wild type (*Pb*A), *app(-)* and *lap(-)* mutants. Ratio of salivary gland versus midgut-derived sporozoites are shown from three independent experiments. **f,g)** Wild type-like range of sporozoite gliding pattern (f) and motility speed (g) of the gene-deletion parasite lines; n: number of sporozoites analysed. **h,i)** Wild type-like range of *in vitro* liver cell infection (j) and liver stage development (k) as evaluated 48 hours post-infection of HepG2 cells. j: three independent experiments with multiple replicates; k: n indicates number of liver stages analysed. **j)** Number of merozoites/per schizont in *in vitro* cultured, mature schizonts of wild type (*Pb*A), *app(-)* and *lap(-)* parasites. n: number of schizonts analysed. **k)** Delayed blood stage development of *app(-)* and *lap(-) mutant* parasite compared to wild type (*Pb*A) as determined from infected red blood cells 26 hours post-invasion of merozoites.

The use of attenuated parasites as vaccines has also been explored in the forms of attenuated blood stages (whole blood stage; Wbs) and sporozoites attenuated in liver stage development (whole sporozoite; Wsp). Attenuation of parasites to generate whole organism vaccines can be achieved by irradiation, application of chemicals or genetic modification ^12–17^. The most advanced Wsp vaccine candidate, the PfSPZ Vaccine, employs sporozoites that have been attenuated by radiation ^18–21^. These sporozoites enter hepatocytes but are unable to replicate and thus abort development early in the liver. The PfSPZ Vaccine, consisting of cryopreserved, vialed radiation-attenuated sporozoites has entered phase 3 clinical development ^15,18,22,23^. Sporozoite attenuation by genetic modification rather than by radiation offers the advantage of a more homogeneous product, increased biosafety for sporozoite production and potentially increased potency ^24^. Several genetically attenuated Wsp vaccines have been developed by the deletion of one to three genes and have been evaluated for inducing protective immune responses in clinical trials ^16,17,25,26^.

The studies using Wsp immunization revealed that Wsp can induce levels of protection that are higher than those achieved with various subunit malaria vaccines, currently under investigation ^1^. In addition to attenuated Wsp, vaccination approaches using Wbs have also been explored ^1,27–29^. However, progress towards a Wbs vaccine is much less advanced compared to Wsp vaccines. Immunization with Wbs has been performed with chemically attenuated ^1,3,28^ and genetically attenuated Wbs ^16,17^ in preclinical studies using rodent malaria models. Immunization with genetically attenuated Wbs of rodent malaria parasites with a reduced growth rate is known to induce robust protection against wild type challenge in mice ^30–34^ but in contrast to the clinically silent Wsp, Wbs immunisation (without also applying drug treatment) harbours the risk of causing a symptomatic blood stage infection as a result of Wbs immunization. Therefore, a high degree of attenuation (and absence of virulence) during blood stage growth is essential. In one study immunization with genetically modified Wbs of *P. falciparum* has been evaluated in humans. In these Wbs the gene has been deleted that encodes the knob-associated histidine-rich protein (KAHRP), which is responsible for the assembly of knob structures at the infected erythrocyte surface. Knobs are required for correct display of the polymorphic adhesion ligand *P. falciparum* erythrocyte membrane protein 1 (PfEMP1), a key virulence determinant. Although the Wbs were immunogenic, administration of higher Wbs doses resulted in blood stage infections with significant parasitemias and malaria-associated symptoms ^29^. Therefore, for further development of safe Wbs vaccines it is important to explore the generation of genetically attenuated Wbs that are avirulent, for example Wbs with a strongly reduced multiplication rate that result in self-resolving infections.

In this study we have screened gene-deletion mutant parasites in the rodent parasite *P. berghei* and the human parasite *P. falciparum* for slow blood stage multiplication rates. In addition, we screened the *P. berghei* mutants for avirulence in mice, i.e. absence of experimental cerebral malaria (ECM), capacity to be transmitted by mosquitoes and for self-resolving blood stage infections. Two of the seventeen screened *P. berghei* mutants showed these characteristics and conferred protection from wild type challenge after single low dose sporozoite immunization. Two of six generated *P. falciparum* gene-deletion mutants showed a slow growth rate. Slow growing, avirulent *P. falciparum* mutants will constitute valuable tools to test induction and breadth of human immune responses and may aid in developing genetically attenuated vaccines to build up liver and blood stage immunity as well as to limit the danger of virulent breakthrough infections of Wsp vaccines.

## Results

### Screening for *P. berghei* gene-deletion mutants with slow blood stage growth for mosquito transmissibility

For creation of mutants with slow growing blood stages we took advantage of the data available from the *Plasmodium* genetic modification project (PlasmoGEM) ^35^. In a large-scale genetic screen ^36^ blood stage growth rates were determined for over 2000 non-clonal gene-deletion mutants. From this screen we selected 50 gene-deletions with reduced growth rate either by 40-60% (set 1) or by 60-80% (set 2). We mostly used the gene-deletion DNA constructs available from the PlasmoGem resource to generate and select clonal mutant parasites for the individual genes. (Figure 1c, figure S1, figure S2, table S1, S2, S3, S4). Of the 50 targeted genes, we were able to select populations of mixed wild type and transgenic parasites for 26 genes (table S1, S2). From those populations we could isolate transgenic clonal lines for 17 genes (Figure 1c). Around half of these 17 clonal mutant parasite lines showed reduced blood stage growth rates comparable to those observed in the PlasmoGEM screen ^36^ with narrow confidence intervals of PlasmoGEM growth rates being a predictor for the growth rate of clonal lines (table S5, figure S3). The other half showed growth rates closer to the growth rate of wild type parasites.

We next analysed life cycle progression of the 17 gene deletion mutants through mosquitoes. Eight mutants showed defects before sporozoite formation. These mutants lacked the genes encoding plasmepsin IV, profilin, beta-catenin like protein 1, telomeric DNA-binding protein (tel DNA-bind protein), mannose-1-phosphate guanyltransferase (man-1-P GT), U2 snRNP-associated SURP motif-containing protein (U2snRNP), diacylglycerol kinase and hypoxanthine-guanine phosphoribosyl transferase (hypox-guan phosph transf) (Figure 1c, table S5). Notably, the other nine mutant lines were able to complete the life cycle (Figure 1c-i, figure S4). These mutants lacked the genes encoding dynein heavy chain, autophagy-related protein 23 (autophagy-rel protein 23), tubulin tyrosine ligase, a conserved protein, dipeptidyl aminopeptidase 1 (dipeptidyl aminopept 1), protein kinase, V-type(+) pyrophosphatase (V-type(+) pyrophosph), aminopeptidase P (app) or M17 leucyl aminopeptidase (lap). The latter two proteins are proteases involved in hemoglobin digestion ^37,38^. In mosquitoes even the two slowest replicating gene-deletion mutants, *app(-)* and *lap(-)*, developed oocysts, albeit at somewhat lower levels than wild type (Figure 1d), resulting in correspondingly lower sporozoite numbers in salivary glands (Figure 1e, table S5). Salivary gland sporozoites of *app(-)* and *lap(-)* mutants showed motility comparable to wild type sporozoites (Figure 1f,g) and infected and developed in *in vitro* cultured hepatocytes similarly as wild type parasites (Figure 1h,i). The reduced blood stage growth of *app(-)* and *lap(-)* parasites ^38^ was not due to a reduced number of merozoites produced per intraerythrocytic cycle (Figure 1j). We noted a prolonged intraerythrocytic developmental time *in vitro* compared to wild type with *lap(-)* parasites were morphologically severely altered and appear immature by giemsa staining. (Figure 1k).

### Blood stage induced infections of five gene-deletion mutants are resolved by mice

To determine virulence (i.e. death or induction of experimental cerebral malaria; ECM) of the 17 gene-deletion mutants, we infected outbred Swiss mice or inbred, highly ECM susceptible C57Bl/6 mice by intravenous injection of 100 blood stages. For 5 out of the 17 mutants infection of C57Bl6 mice led to a delayed infection that could be controlled and cleared by the mice. These 5 mutants lacked the genes encoding lap, app, U2 snRNP, diacylglycerol kinase and hypoxanthine-guanine phosphoribosyl transferase (Figure 1c, Figure 2a-c, figure S5-S7, table S6). As illustrated for *app(-)* or *lap(-)* blood stage-induced infections of C57Bl/6 mice in Figure 2a-c, the infection peaked around day 14 and 15, respectively and was ultimately resolved within 20 to 21 days (Figure 2a-c, Table 1, table S7). Infection of Swiss mice led to a delayed infection in 4 out of the 17 mutants that could be controlled and cleared. These mutants lacked the genes encoding lap, app, U2 snRNP and hypoxanthine-guanine phosphoribosyl transferase (Figure 1c, Figure 2a-c, figure S5-S7, table S6). In Swiss mice infected with *app(-)* or *lap(-)* blood stages, parasitemia peaked on average on day 18 and was resolved by day 21 (Table 1, table S7). Only one Swiss mouse infected by *app(-)* blood stages died due to anemia at day 21 post-infection. Infections with wild type or the other gene-deletion mutants were not self-resolving and resulted in death of part or all animals (Figure 1c, figure S5-S7, table S6).

**Fig. 2.**
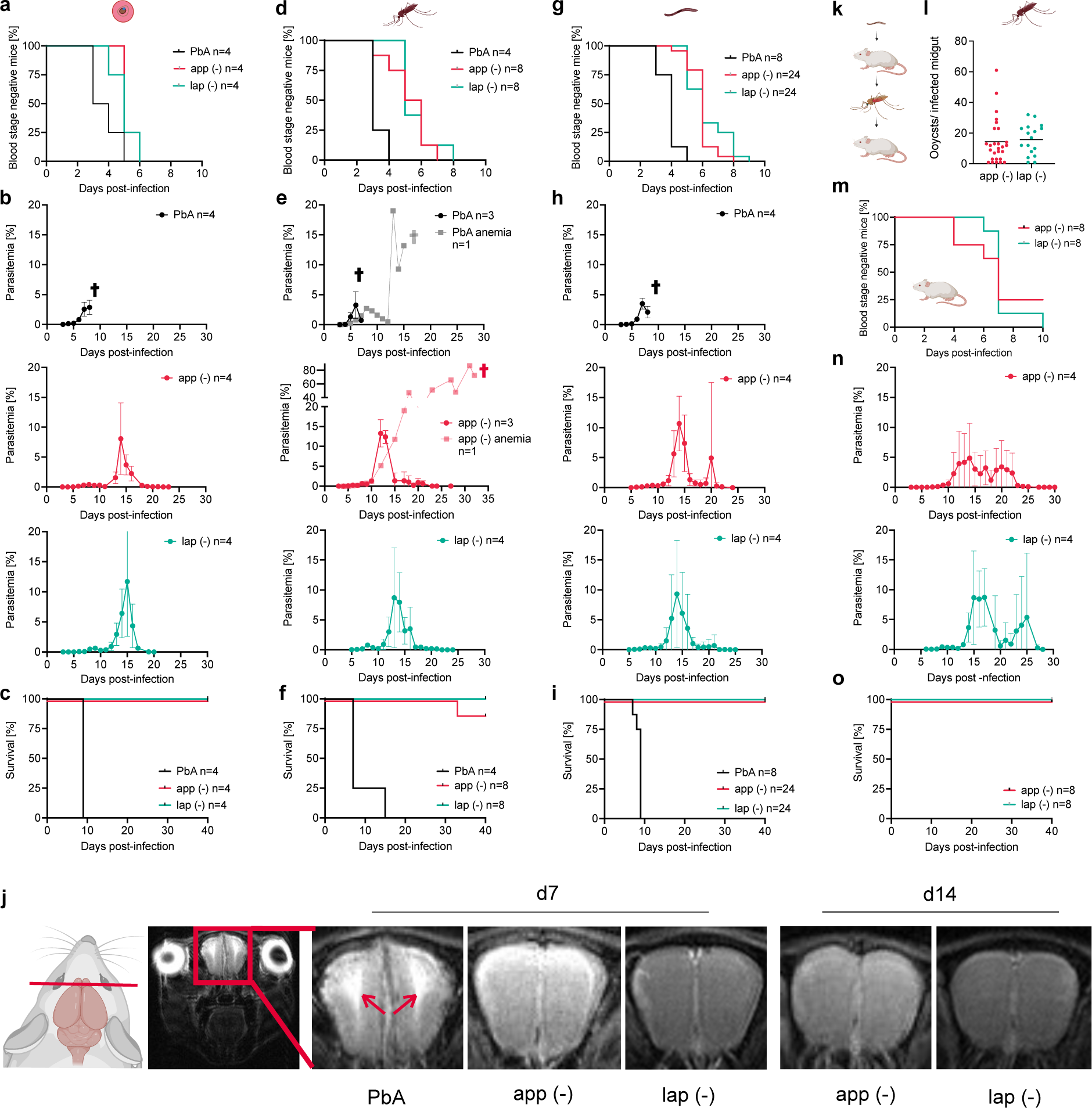
Sporozoites lacking *app* and *lap* induce self-limiting blood stage infections. **a)** Percentage of blood stage negative C57Bl/6 mice infected with 100 wild type, *app(-)* and *lap(-)* blood stage parasites. n: numbers of infected mice. **b,c)** Parasitemia (b) and mouse survival (c) for the animals infected in a. Mean parasitemia values +/− standard deviation are shown. **d)** Percentage of blood stage negative C57Bl/6 mice infected with wild type, *app(-)* and *lap(-)* sporozoites by 10 biting mosquitoes. n: numbers of infected mice. **e,f)** Parasitemia (e) and mouse survival (f) for the animals in d. Light grey parasitemia curve shows the infection course of the *app(-)* infected mouse that died. Mean parasitemia values +/− standard deviation are shown. **g)** Percentage of blood stage negative C57Bl/6 mice infected with 1.000 wild type, *app(-)* and *lap(-)* salivary gland-derived sporozoites. n: numbers of infected mice. **h,i)** Parasitemia (h) of one experiment with four mice and (i) mouse survival for animals infected in g. Mean parasitemia values +/− standard deviation are shown. **j)** MRI images showing brain oedema at the level of the olfactory bulb in mice infected by wild type (*Pb*A) at 7 days post infection with 1.000 sporozoites but not in mice infected with *app(-)* and *lap(-)* sporozoites. **k)** *app(-)* and *lap(-)* parasites can re-transmit to mosquitoes (cartoon). Oocyst prevalence is shown for mosquitoes that were fed on mice 14 days after the mice were infected with *app(-)* and *lap(-)* sporozoites. **l)** Prepatency of C57Bl/6 mice infected by caged mosquitoes carrying re-transmitted *app(-)* and *lap(-)* parasites (mosquitoes from experiment described in k). 6 out of 8 *app(-)* and all *lap(-)* infected mice became blood stage patent. **m, n)** Course of blood stage infection (m) and survival (n) of mice from l.

**Table 1.** Summary of wild type, *app(-)* and *lap(-)* infections of Swiss and C57Bl/6 mice. **a)** Blood stage-initiated infections by intravenous injection of 100 infected red blood cells (iRBC). **b)** Mosquito stage-initiated infections by the bite of 10 mosquitoes per mouse (natural transmission) or the intravenous injection of 1.000 or 10.000 sporozoites. Numbers for peak parasitemia percentage and days are averages from the indicated number of investigated mice.

|  | 100 iRBC Swiss |  |  | 100 iRBC C57Bl/6 |  |  |
| --- | --- | --- | --- | --- | --- | --- |
|  | Mice clearing/<br>mice infected<br>(prepatency [d]) | Peak parasitemia<br>[%] (d) | Blood stage cleared<br>(d) | Mice clearing/<br>mice infected<br>(prepatency [d]) | Peak parasitemia<br>[%] (d) | Blood stage cleared<br>(d) |
| PbA | 0/3 (6) | 27 | n.a. | 0/4 (4) | 4 | n.a. |
| lap (-) | 4/4 (6) | 17 (18) | 21 | 4/4 (5) | 11 (15) | 20 |
| app (-) | 3/4 (5) | 22 (18) | 21 | 4/4 (6) | 9 (14) | 21 |

b)
|  | Natural transmission<br>C57Bl/6 |  |  | 1.000 salivary gland sporozoites i.v.<br>C57Bl/6 |  |  | 10.000 salivary gland sporozoites i.v.<br>Swiss |  |  |
| --- | --- | --- | --- | --- | --- | --- | --- | --- | --- |
|  | Mice clearing/<br>mice infected<br>(prepatency [d]) | Peak parasitemia<br>[%] (d) | Blood stage cleared<br>(d) | Mice clearing/<br>mice infected<br>(prepatency [d]) | Peak parasitemia<br>[%] (d) | Blood stage cleared<br>(d) | Mice clearing/<br>mice infected<br>(prepatency [d]) | Peak parasitemia<br>[%] (d) | Blood stage cleared<br>(d) |
| PbA | 0/4 (3) | 6 | n.a. | 0/8 (4) | 4 | n.a. | 0/6 (5) | 23 | n.a. |
| lap (-) | 8/8 (6) | 16 (15) | 22 | 24/24 (6) | 19 (17) | 23 | 4/4 (7) | 7 (16) | 20 |
| app (-) | 7/8 (5) | 12 (14) | 22 | 24/24 (6) | 19 (17) | 27 | 6/6 (7) | 14 (16) | 23 |

n.a. not applicable

### Sporozoite induced blood stage infections with *app(-)* and *lap(-)* are resolved by mice

Next we assessed the virulence of sporozoite-induced infections of the gene-deletion mutants, *app(-)* and *lap(-)*, which have already shown full clearance in C57Bl/6 mice after iRBC-induced infections and were able to produce infectious sporozoites after mosquito infection (Figure 1c, Figure 2, table S6, table S7). Infection of C57Bl/6 mice by bite of infected mosquitoes or by intravenous injection of 1.000 *app(-)* or *lap(-)* sporozoites or Swiss mice infected by intravenous injection of 10.000 *app(-)* or *lap(-)* sporozoites resulted in blood stage infections but with a significantly (p<0.0001, Dunnett’s multiple comparisons test) reduced blood stage growth rate compared to mice infected with wild type sporozoites as assessed at day 6 post-infection (Table 1, Figure 2d-I, figure S8). Sporozoite-induced wild type infections resulted in death of all 16 mice (Figure 2f,l, Table 1, figure S8). In contrast, out of 38 *app(-)* infected mice only one C57Bl/6 mouse infected with *app(-)* sporozoites died (Figure 2f,i, Table 1, figure S8). All 36 mice infected with *lap(-)* sporozoites survived (Figure 2f,i, Table 1, figure S8). In mice surviving from a sporozoite-induced infection, the infection peaked around day 14 to 17 and was ultimately resolved within 20-27 days (Table 1, table S7). The highest measured parasitemia was 70% (C57Bl/6 infected with 1.000 *app(-)* sporozoites) and the longest time to complete parasite clearance was 46 days with mice being infected for 39 days (C57Bl/6 infected with 1.000 *app(-)* sporozoites; table S7). Even at high parasitemias all surviving *app(-)* and *lap(-)* infected mice showed no signs of disease. To evaluate possible signs of ECM, C57Bl/6 mice were subjected to magnetic resonance imaging (MRI) on day 7 and 14 post-infection with 1.000 salivary gland sporozoites to assess brain morphology and appearance of edema ^39^. While mice infected with wild type parasites showed clear signs of ECM already at low parasitemias at day 7 post-infection, no signs of ECM were detected in *app(-)* or *lap(-)* infected mice (Figure 2j). Even when *app(-)* and *lap(-)* infected mice showed high parasitemias around day 14 post-infection, no signs of edemas in the olfactory bulb were detectable. Feeding of mosquitoes on mice infected with *app(-)* and *lap(-)* sporozoites prior to resolving the blood stage infection resulted in infection (Figure 2k,l). These mosquitoes were able to infect C57Bl/6 mice 21 days post-infection demonstrating that these mutant parasites can be cycled between vector and host (Figure 2m-o).

### Mice infected with *app(-)* and *lap(-)* parasites are protected from challenge with wild type sporozoites

We next investigated if C57Bl/6 mice that cleared an *app(-)* or *lap(-)* infection could survive a challenge with wild type sporozoites. To this end we challenged mice that resolved a *app(-)* and *lap(-)* blood stage- or sporozoite-induced infection and remained parasite free for at least 60 days with wild type sporozoites at 90, 180 and 360 days post-infection (Figure 3a-h, Table 2, table S8). Around half of the challenged mice did not develop a detectable blood stage infection after challenge at 90 days (Figure 3b, Table 2, table S8). The other half developed a low-level blood stage infection with the onset of all blood stage infections delayed by 3 to 9 days compared to age-matched naïve mice. The blood stage infections peaked between 9 and 19 days at less than 2% parasitemia and were cleared after approximately 20 days (Figure 3; Table 2, figure S9). When challenged at 180 days, 60% (*app(-)*) and 20% (*lap(-)*) of the mice did not develop a blood stage infection. The mice that developed a blood stage infection showed parasitemias of maximum 2.4% for *lap(-)* and the longest time to complete parasite clearance was 19 days with mice being infected for 13 days. Challenge after 360 days resulted in blood stage infections of 4 out of 4 mice (*app(-),* average peak parasitemia of 2.3%), and 1 out of 4 mice (*lap(-),* peak parasitemia of 0.7%) (Table 2, table S8). All challenged mice were able to resolve the infection and survived.

**Fig. 3.**
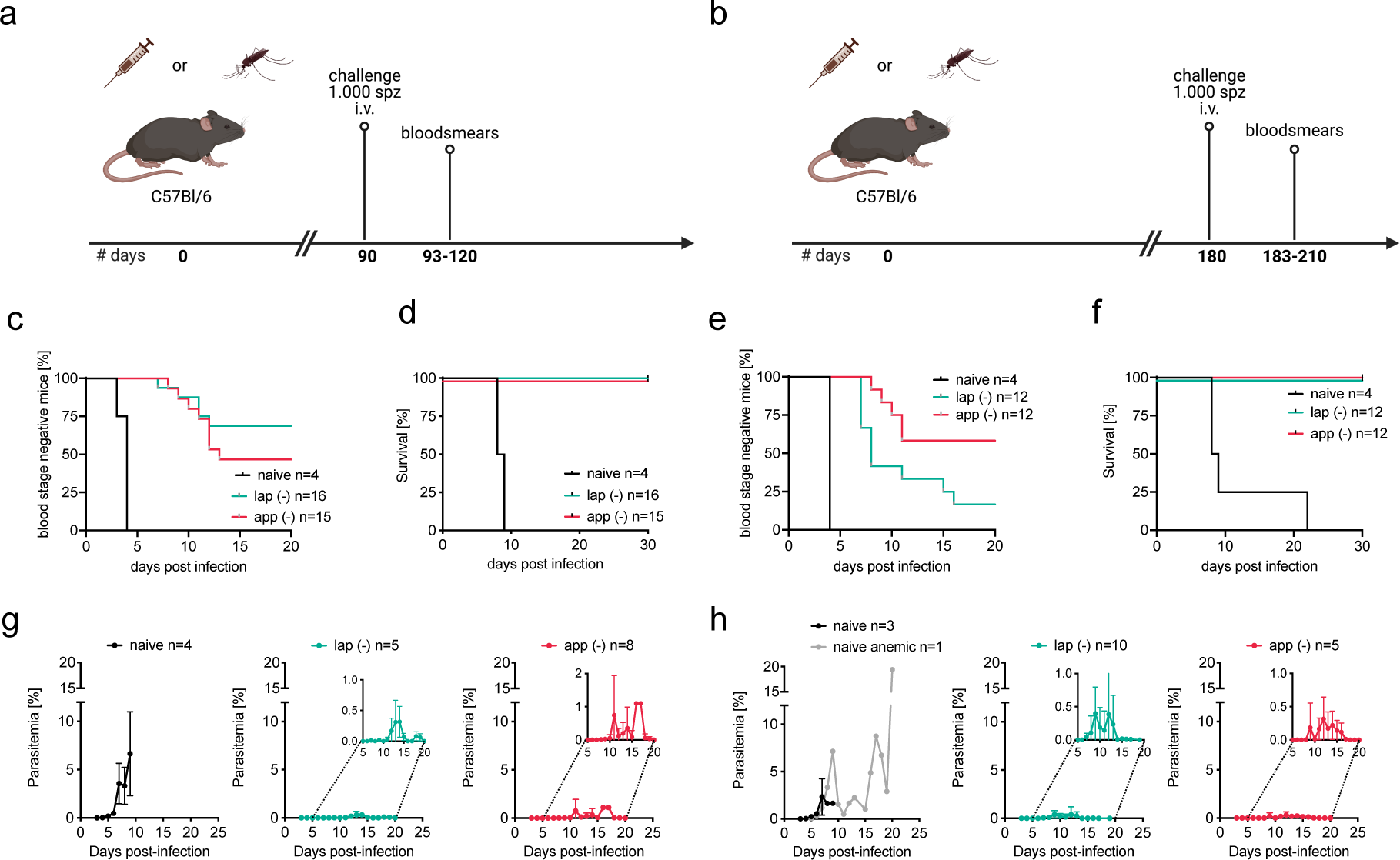
Self-limiting infections cause protection from lethal wild type challenge. **a,b)** Illustration of challenge experiments. C57Bl/6 mice were infected with 1.000 salivary gland-derived wild type sporozoites 90 (a) or 180 (b) days post immunization with either 1.000 salivary gland-derived sporozoites or by natural transmission of *app(-)* or *lap(-)* parasites. **c-f)** Percentage of blood stage negative (c,e) and surviving (d,f) mice after challenge with wild type sporozoites after 90 (c,d) and 180 (e,f) days. **g,h)** Parasitemia of naïve or immunized C57Bl/6 as challenged in a or b, respectively. Small insets display magnifications of parasitemia curves.

**Table 2.**
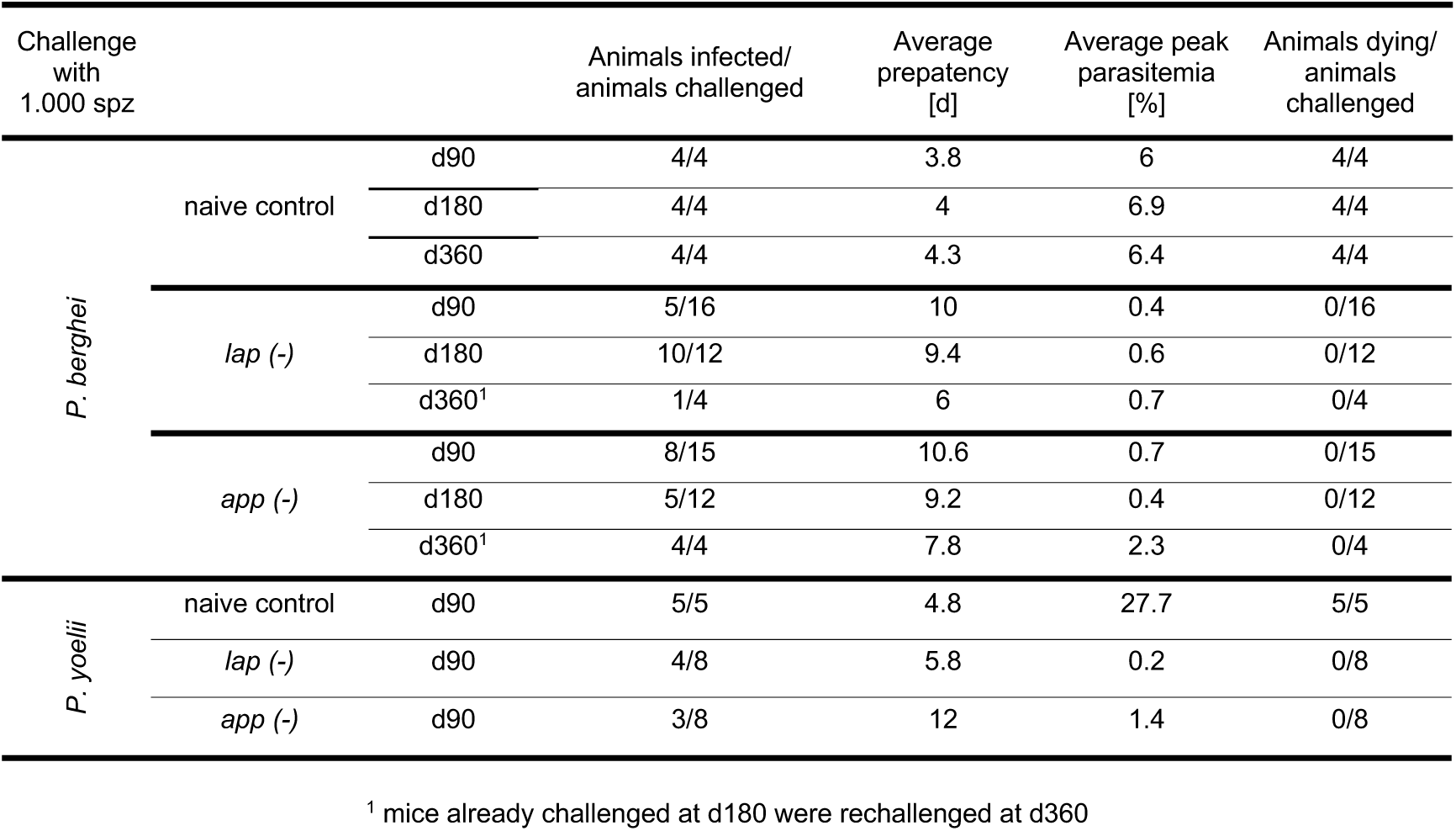
Protection of C57Bl/6 mice immunized with *app(-)* or *lap(-)* sporozoites against wild type challenge 90 days, 180 days or 360 days post immunization.

Finally, we tested if mice immunized with either *app(-)* or *lap(-) P. berghei* sporozoites that resolved infections were protected from heterologous challenge. To this end we challenged the mice with 1.000 *P. yoelii* sporozoites 90 days post-infection. Only half of the mice developed a *P. yoelii* blood stage infection. In contrast to infected naïve control mice, which all died, all *app(-)* or *lap(-)* immunized mice survived this heterologous challenge, indicating cross-species immunity (Table 2). *P. yoelii* induced infections appeared as fast in the blood as *P. berghei* in the *app(-)* immunized mice but showed higher peak parasitemias than in *P. berghei* challenges (Table 2). However, *P. yoelii* induced infections in *lap(-)*-immunized mice appeared 4 days earlier than when mice were challenged with *P. berghei* sporozoites (Table 2). Intriguingly, the *P. yoelii* challenged mice showed a lower average peak parasitemia (0.2%) than *P. berghei* challenged mice (0.4%).

Together, these results suggest that differences exist between *app(-)* and *lap(-)* parasites in inducing immunity by infection with sporozoites. Importantly, our data show that a single infection with low sporozoite numbers of gene-deletion mutants with reduced blood stage growth and virulence can result in complete in protection against challenge with wild type parasites.

### Generation of slow-growing P. falciparum mutant parasites

To generate *P. falciparum* gene deletion mutants with a slow blood-stage growth rate we used CRISPR/Cas9-based methods to delete (parts of) selected genes and introduce premature stop codons. In addition, as control mutants we introduced silent mutations into the selected/target genes (Figure 4, figure S10a,b), performing in total over 50 independent transfections. We first targeted the *P. falciparum app* and *lap* genes for deletion. These two *P. falciparum* genes were previously reported to be refractory to gene deletion, indicating they are essential for *in vitro* blood stage growth *(37, 39)*. In line with these observations, we were not able to delete these *P. falciparum* genes in multiple transfection attempts (table S9, figure S10c,d). We next targeted an additional six genes based on evidence for slow growth of gene-deletion mutants obtained in genome-wide knockout screens in *P. berghei* and *P. falciparum* and published reports (table S9 ^40–44)^. These six genes encode for *Plasmodium* exported protein (HYP11), thioredoxin 2 (TRX2), TIF eIF-2B delta subunit (EIF-2B), nicotinamide mono nucleotide adenylyl transferase (NMNAT), serine/ threonine protein kinase FIKK family (FIKK8), and mago nashi protein homologue (Mago Nashi). We generated and selected gene-deletion mutants and matching control lines with silent mutations for all these six genes (Figure 4, figure S10e-j). For two mutants we observed a slow proliferation of their blood stages *in vitro* at 80% and 70% of the control growth rate (Figure 4c,d). These mutants either lack the gene encoding TRX2 or FIKK8, respectively. The other four mutants showed a wild type like asexual growth rate (Figure 4c,d, table S9). Analysis of *Pf trx2(-)* blood stage development in synchronized cultures revealed a delay in the development of late trophozoites/early schizonts, similar to the reported delay in trophozoite/schizont maturation of *P. berghei trx2(-)* mutants ^43,44^ (Figure 4e). These results show that it is also possible to generate and select clones of slow growing blood stages of *P. falciparum* gene-deletion mutants, despite the lower transfection efficacy and longer asexual cycle of *P. falciparum* compared to *P. berghei*. Importantly, the CRISPR-Cas9-based approach used in this study results in generation and selection of gene-deletion mutant lines that lack residual foreign genetic material used to delete the target genes, for example drug-selectable markers, or sequences of the CRISPR/Cas9 constructs. The absence of such DNA is essential for approval of future use of these mutant parasites as genetically attenuated Wbs in immunization approaches in humans.

**Fig.4.**
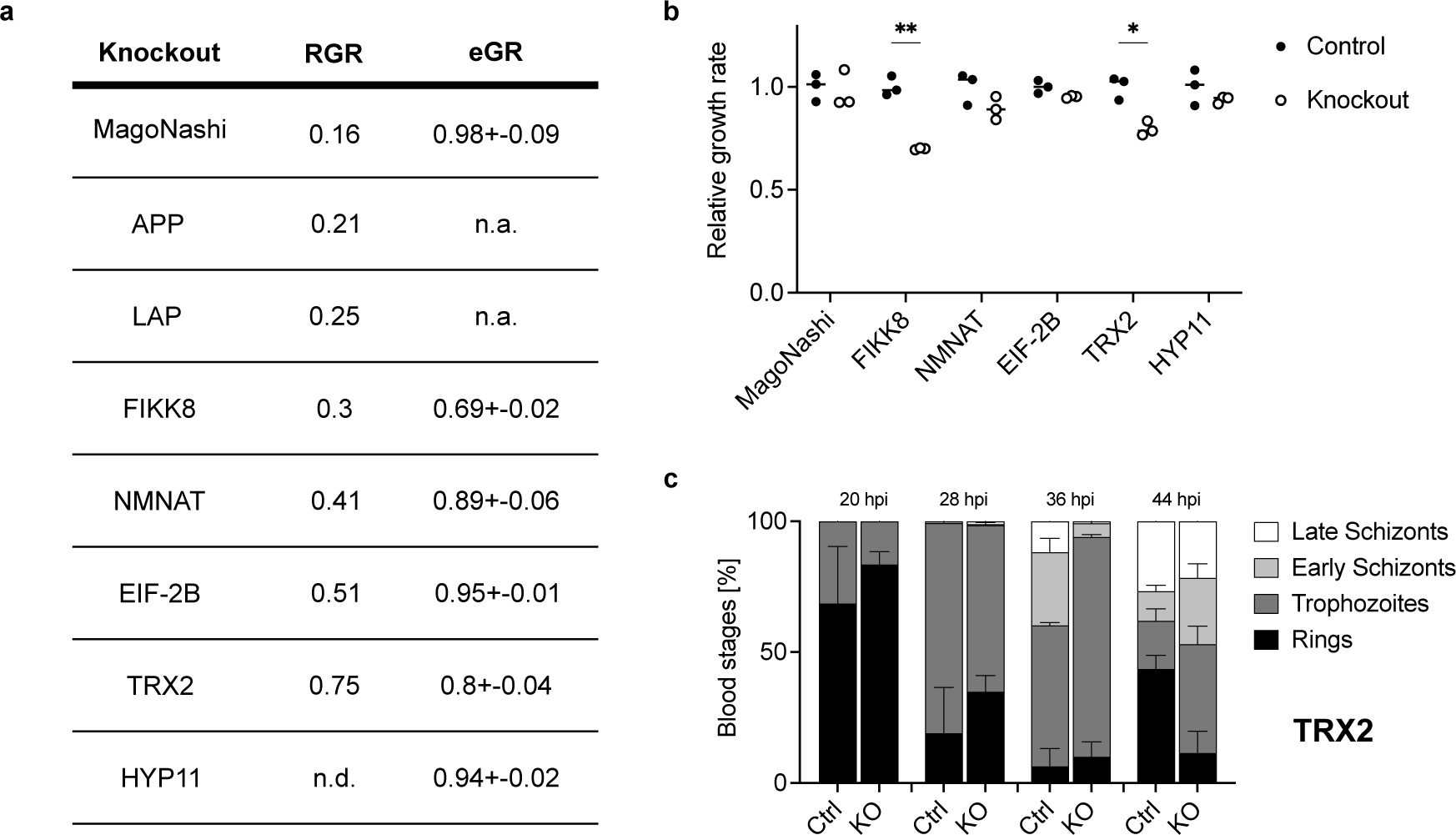
Generation of slow-growing *P. falciparum* gene deletion lines. **(a,b)** Predicted (RGR; ^36^ and experimental (eGR; this study) growth rates of parasite lines; for gene ID, please see Supplemental Table 9. Per knockout, growth rates were normalised to the mean of the respective control line that was generated by introduction of silent mutations. Unpaired t-test with Holm-Sídák‘s correction for multiple comparisons, * p < 0.05, ** p < 0.01. (**c**) Stage composition of synchronised *Pf trx(-)* and matching control lines. 100 parasites were staged per replicate and time point. At least three independent experiments were conducted. RGR, relative growth rate; eGR, experimental growth rate; hpi, hours post infection; Ctrl, control; KO, knockout.

## Discussion

Here we screened seventeen new *P. berghei* gene-deletion mutants with attenuated blood stage growth for use as experimental whole blood stage (Wbs) and whole sporozoite (Wsp) immunization approaches. We identified two mutants, *lap(-)* and *app(-)*, that could complete the parasite life cycle and showed slow blood stage growth in mice both after infection with sporozoites and with blood stage parasites. All 44 mice infected with either *lap(-)* blood-stages or sporozoites resolved blood-stage infections within less than 4 weeks and remained parasite free (analysed by Giemsa stained blood smears) for up to 180 days. This corresponds to a 95% binomial confidence interval of 0-8%. Following challenge with wild type *P. berghei* or *P. yoelii* sporozoites these mice were either sterilely protected or cleared blood stage infections without showing signs of disease. *App(-)* parasites showed similar characteristics although 2 out of 46 mice infected with blood-stages or sporozoites were not able to resolve the blood-stage infection and died. This higher pathogenicity of *app(-)* compared to *lap(-)* correlates with a more severe growth attenuation of *lap(-)* blood stages *in vivo* and *in vitro*.

The generation of *P. falciparum* parasite lines with similar strongly reduced growth and virulence as for *P. berghei lap(-)* blood stages will help to establish controlled human infections with longer exposure of human volunteers to blood stage infections before clearance with antimalarials. This will permit to gain novel insights into immune responses induced by genetically attenuated Wbs in humans. However, we and others were not able to produce similar *P. falciparum* mutants lacking the gene encoding M17 leucyl aminopeptidase using a CRISPR/Cas9 gene-deletion approach ^37,45^, excluding further analysis on both growth and virulence features of such *P. falciparum* Wbs. We therefore continued to generate and select *P. falciparum* mutants with a slow growth rate selecting target genes from previously published studies ^36,41,43,44^. In a small-scale gene-deletion screen we show that it is also possible to generate and select slow growing blood stages of *P. falciparum* gene-deletion mutants, despite the lower transfection efficacy and longer asexual cycle of *P. falciparum* (48 h) compared to *P. berghei* (24 h).

Recently, a genetically attenuated Wbs vaccine was tested with the candidate line lacking the knob-associated histidine rich protein KAHRP, which is involved in adhesion of infected red cells to the endothelium ^29^. This Wbs vaccine leads to increased splenic clearance of iRBCs and hence lowered virulence. However, high doses of parasites led to patent infections ^29^. Additional deletion in this *kahrp(-)* parasite line of a gene from a growth retarded parasite line as presented here might allow the use of higher parasite numbers for immunization and lead to longer infections in humans. Additionally, combination of gene-deletions resulting in late liver stage attenuated parasite lines ^16,17,25,46^ with mosquito-transmittable blood stage attenuated parasites could safeguard from virulent breakthrough infections.

In rodent malaria parasites many other candidate genes could be investigated for their potential to cause self-limiting infections once ablated from the genome. Bushell et al. identified 256 *P. berghei* genes that bestow a relative growth rate of 0.2-0.4 and another 133 with a relative growth rate of 0.4-0.6 upon their deletion ^36^. As we were able to identify 2 candidates out of our list of 50 targeted genes, simple interpolation would posit that from the remaining 342 genes around 15 more parasite lines with similar characteristics should be obtainable. However, growth rates of our individual gene deletion mutants of both *P. berghei* and *P. falciparum* showed only a limited overlap with predicted growth rates of the genome wide screens ^36,41^.

Our study also provides the basis for several additional strategies to generate attenuated blood stage parasites with reduced growth and virulence that produce sporozoites and thus are transmissible. For example, when gene deletion leads to reduced growth and/or virulence of blood stages but in addition blocks further development in the mosquito, promoter swap strategies could be employed to express the gene during mosquito development by expressing the respective gene from a gametocyte or mosquito stage specific promoter ^47–50^. In addition, mutant parasites with slow growing blood stages but still showing virulence characteristics could be used as a basis to generate double or triple gene-deletion mutants that might attenuate virulence. Finally, the human-infecting zoonotic *P. knowlesi* parasite could also be tested for slow growth as an intermediate pre-clinical model ^51,52^ as it can be cultured *in vitro* like *P. falciparum* but shows a higher transfection efficacy and a fast red blood stage cycle of just 24 hours ^53^. Lastly, slow growing *P. falciparum* parasites were recently also identified by a screen using selection linked resistance marker integration, which could help to prescreen for gene-deletions resulting in an avirulent phenotype ^54^.

In conclusion, we screened a number of parasite gene-deletion mutants to obtain blood stage attenuated malaria parasites in *P. berghei* and *P. falciparum*. We generated mutant *P. berghei* lines with reduced growth of blood stages and infection of mice by a low number of infected-mosquito bites or by single injection of sporozoites of these mutants led to protection from disease after challenge with wild type sporozoites. This new type of Wbs-attenuation will constitute valuable tools to inform on the induction of immune responses and may aid in developing as well as safeguarding attenuated parasite vaccines.

## Acknowledgements

We thank the PlasmoGEM consortium for generously supplying the transfection vectors for this project; Miriam Reinig for mosquito breeding; Steffen Borrmann, Elias Farr, Kai Matuschewski, Katja Müller and Mirko Singer for helpful discussions and suggestions regarding the manuscript. We thank specifically the late Shahid Khan for motivating and lively discussions. Figures were created partially with BioRender.com.

## Methods and Materials

### Bioinformatic identification of slow growing parasites

Gene-deletion mutants showing a growth rate of 0.2 – 0.6 were retrieved from PlasmoGEM (https://plasmogem.umu.se/pbgem/). In addition, from this list of genes/mutants, genes were selected either manually or using PlasmoSPOT (https://frischknechtlab.shinyapps.io/plasmoSPOT/) ^55^ for a transcription profile that shows transcription in asexual blood stages but low or no expression in gametocytes and mosquito stages.

### Ethics statement

All experiments were performed in accordance with GV-SOLAS and FELASA guidelines and have been approved by the German authorities (Regierungspräsidium Karlsruhe; G283/14, G-8/21, G9/21).

### Animals, parasite and mosquito

SWISS mice were obtained from Janvier Laboratories (minimum 20 g) and C57Bl/6 from Charles River laboratories (minimum 18g). *Plasmodium berghei* ANKA, *P. yoelii* 17XL and *Anopheles stephensi* were used for all experiments.

### Generation of *P. berghei* gene-deletion mutants

Gene-deletion DNA constructs containing the hDHFR/ yFCU selectable marker cassette, allowing selection of transgenic parasites by pyrimethamine treatment of mice, were obtained from PlasmoGem ^35^. Transfections were performed as described previously using synchronized *P. berghei* ANKA schizonts cultured overnight in RPMI-1640 complete (RPMI-1640 (GIBCO) supplemented with 20% FBS (US origin GIBCO) and 0.03% Gentamycin (10 mg/ml; PAA, Pasching, Austria)) at 37°C, 5% CO_2_ ^56^. Briefly, purification of cultured schizonts was performed through a 55% Nycodenz/ PBS gradient (Axis-shield diagnostics, Heidelberg, Germany). 10 µg plasmid was digested with NotI overnight at 37°C followed by ethanol precipitation and resuspension in 20 µl PBS. The linearized construct and the purified schizonts were mixed with Nucleofector solution from an Amaxa human T cell Nucleofector Kit and electroporated using the Amaxa Nucleofector II device (Lonza, Köln, Germany). Directly after transfection 50 µl RPMI-1640 complete was added to the transfection reaction followed by intravenous injection into one SWISS mouse. 24 hours post-transfection drinking water was exchanged to 280 µM pyrimethamine (Sigma-Aldrich, Munich, Germany) to select for transgenic parasites. Parasitemia was assessed from day 7 onwards through Giemsa-stained thin blood smears. Once parasitemia reached 1.5% the blood was collected through cardiac puncture using a 1 ml syringe containing 50 µl heparin and mixed for storage in liquid nitrogen as follows: 100 µl blood, 200 µl Freezing solution (Alsever solution (Sigma-Aldrich, Munich, Germany) + 10% glycerol. In addition, genomic DNA was isolated from iRBCs by saponin-lysis and followed by using the DNeasy Blood & Tissue Kit (Qiagen). Upon confirmation of correct integration by PCR isogenic parasite lines were generated by limiting dilution in mice ^57^. Only mutant lines with correct integration of the gene-deletion constructs and absence of the wild type gene were used for further analysis. Primers used for genotyping PCRs are shown in table S3.

### Blood stage and mosquito infection

Blood was taken by cardiac puncture from a donor mouse at 0.2-1% parasitemia and upon serial dilution in PBS 100 infected red blood cells (per 100 µl PBS) were injected intravenously. Prepatency as well as parasitemia was assessed from day three post-infection.

For mosquito infections, blood was taken by cardiac puncture from a donor mouse at 1-3% parasitemia. Recipient SWISS mice were treated with phenylhydrazine (0.6% in PBS, Sigma-Aldrich, Munich, Germany) to induce reticulocytes three days prior to being infected with 20 million iRBCs. 3 days after infection exflagellation was assessed and starved mosquitoes were allowed to bite anesthetized mice for 20 minutes.

### Evaluating mosquito infections

Presence of midgut oocysts was evaluated 10-14 days post-infection by first permeabilizing midguts with 0.01% NP40/PBS for 30 min followed by staining with 0.1 % mercurochrome/H_2_O for 1 hour. Following three washing steps with PBS, stained midguts were mounted on a glass slide, covered with a coverslip and assessed using a Zeiss Axiovert 200 M inverted microscope with a 10x objective.

Number of sporozoites was assessed 18 days post-infection by isolating midguts and salivary glands of 10 mosquitoes. Sporozoites were released into RPMI-1640 by mechanically disrupting the tissue within 1.5 ml reaction tubes using a pestle and counted in a Neubauer chamber on a Zeiss Axiostar light microscope under a 40x phase contrast objective.

*In vitro* motility assays were performed essentially as described before ^58^ with salivary gland derived sporozoites isolated between day 18-24 post-infection and purified with an Accudenz density gradient ^59^. Imaging was performed on a Zeiss Axiovert 200 M fluorescent microscope (25x objective, DIC, frame rate of 1/3s for 3 minutes). FIJI ^60^ was used for analysis of motility patterns as well as for calculation of sporozoite speed.

### *In vivo* infections

For natural transmission experiments 10 mosquitoes were transferred to a small cup one day before the experiment and starved overnight. The next day C57Bl/6 were anesthetized and placed on top of the cups. Mosquitoes were allowed to feed for 10 minutes. Alternatively, sporozoites were isolated 18-23 days post-infection from mosquito salivary glands, sporozoites released, counted and injected into SWISS (10.000 sporozoites) or C57Bl/6 (1.000 sporozoites) mice by intravenous inoculation into the tail vein. Prepatency and course of parasitemia was assessed from day 3 post-infection onwards by Giemsa-stained smears from peripheral blood.

### Magnetic resonance imaging

C57Bl/6 were infected by intravenous injection of 1.000 salivary gland derived sporozoites and parasitemia was monitored by Giemsa-stained blood smears starting day 3 post-infection. From day 6 post-infection MRI of mice was performed with a 9.4-T horizontal bore small-animal MRI unit (BioSpec 94/20 USR; Bruker BioSpin, Ettlingen, Germany) with a four-channel phased-array surface receiver coil as described before ^61^. Mice were anesthetized with 2% isoflurane. Anesthesia was maintained with 0.5%–1.5% isoflurane. Animals were kept on a heating pad to keep the body temperature constant, and respiration was monitored externally during imaging with a breathing surface pad controlled by an in-house–developed LabView program (National Instruments, Munich, Germany). To exlude the earliest manifestion of the experimental malaria, the olflactory bulb was imaged with a 2D T2-weighted sequence (TR/TE = 2,000/22 ms, slices = 12, and slice thickness = 0.7 mm).

### Challenge infections

Mice immunized with either 100 iRBC, 1.000 infectious sporozoites or by mosquito bite were rechallenged 90 days, 180 days or 360 days after infection using 1.000 freshly dissected PbA wildtype sporozoites. Appearance of blood stage parasites was monitored daily starting three days post-challenge by Giemsa-stained bloodsmears for at least 20 days. Mice challenged at 360 days received one previous challenge infection at d180.

### *In vitro P. berghei* hepatocyte invasion and exo-erythrocytic development

Hepatocyte invasion and exoerythrocytic development was performed in HepG2 cells cultivated at 37°C, 5% CO_2_ in DMEM complete (10% FCS, 1% Penicillin/ Steptomycin; Gibco). Two days prior to infection with 10.000 sporozoites, 100.000 (invasion) or 30.000 cells (exo-erythrocytic development) were seeded per well of an 8-well Permanox Lab-Tek chamber slide (Nunc). After two hours of incubation with sporozoites at 37°C, 5% CO_2_ wells were washed twice with DMEM complete and the invasion assay was stopped by adding 4% PFA for 20 min at RT while fresh DMEM complete supplemented with an Antibiotic-Antimycotic cocktail (1x, Gibco) was added to the other wells. After 48 h incubation at 37°C, 5% CO_2_ cells were fixed by adding ice cold methanol for 10 min at RT. Both assays were washed twice with 1% FCS/ PBS and then blocked for 2 h at 37°C or o/n 4°C in 10 % FCS/ PBS.

Cells in the invasion assay were stained subsequently for 1 h at 37°C with primary mouse αCSP antibodies (cell culture supernatant 1:300 diluted in 10% FCS/ PBS) and anti-mouse Alexa Fluor 488 antibodies (1:300 diluted in 10% FCS/ PBS; Life technologies). Following two washing steps with 1% FCS/PBS the cells were permeabilized by addition of ice-cold methanol for 10 min at RT. Subsequently another round of blocking with 10% FBS/ PBS at 37°C or at 4°C overnight was followed by a second round of subsequent incubation with αCSP (cell culture supernatant 1:300 diluted in 10% FCS/ PBS) and secondary anti-mouse Alexa Fluor 546 antibodies (1:300 diluted in 10% FCS/ PBS; Life technologies) for 1 h at 37°C. Hoechst 33343 was added (1 µg/ ml; Life technologies) for 5 min at RT to visualize DNA. After two washing steps using 1% FCS/ PBS the assays were mounted.

Cells in the exo-erythrocytic development assays were subsequently stained with a primary antibody against *Plasmodium berghei* HSP70 (cell culture supernatant 1:300 diluted in 10% FCS/ PBS) and an anti-mouse Alexa Fluor 488 (1:300 diluted in 10% FCS/ PBS; Life technologies). Hoechst 33343 was added after for 5 min at RT and the cells washed twice in 1% FCS/ PBS the assays before being mounted. Cells were assessed at a Zeiss Axiovert 200 M inverted microscope and Fiji was used to measure sizes of liver stages.

### *In vitro P. berghei* blood stage development

Blood of infected mice was collected after mosquito infection, transferred to a 75 cm^2^ cell culture flask containing RPMI-1640 supplemented with 20% FCS and 0.03% gentamycin and cultivated at 37°C, 5% CO_2_. 16 hours later schizonts were purified using a 55% Nycodenz gradient and centrifugation at 209 g for 25 min without break. Schizonts were collected from the interphase and washed once with RPMI-1640 supplemented with 20% FCS and 0.03% gentamycin. Purified schizonts were intravenously injected into a naïve Swiss. Two hours after infection blood was collected via cardiac puncture and transferred to a 75 cm^2^ cell culture flask containing RPMI-1640 supplemented with 20% FCS and 0.03% gentamycin and cultivated at 37°C, 5% CO_2_. Starting 24 h post-infection and then every two hours 1 ml of culture was collected, and development of blood stages was evaluated via Giemsa-stained blood smears. Developmental stages were classified as trophozoite (one nucleus), early schizont (multiple nuclei visible) or late schizont (single merozoites visible).

### Cloning of constructs for the generation of *P. falciparum* knockout lines

To generate genetic deletion lines of target genes, we used a CRISPR/Cas9-based genetic strategy resulting in the loss of several hundred base pairs of gene sequence as well as the introduction of a pre-mature stop codon and a frame shift (figure S9a). In parallel, control lines using the same gRNA only introduced a silent shield mutation into the target gene (figure S9b). Plasmids were based on pDC2-cam-coCas9-U6-hDHFR (a gift from M. Lee ^62^). We designed gRNAs using the Eukaryotic Pathogen CRISPR gRNA Design Tool ^63^ and ordered forward and reverse sequence as unmodified single-stranded oligonucleotides (Merck). After annealing of the forward and reverse oligonucleotide, these were ligated into the BbsI-digested pDC2-cam-coCas9-U6-hDHFR plasmid. Homology regions were amplified by PCR using the primers indicated in table S3 and cloned into the EcoRI/AatII-digested vector by Gibson assembly (NEBuilder HiFi DNA Assembly, New England Biolabs GmbH). Plasmid sequences were verified by control digest and sequencing. For transfection, medium-scale plasmid preparations were produced using the NucleoBond Xtra Midi Kit (Macherey-Nagel).

### *Plasmodium falciparum* culture and transfection

*P. falciparum* 3D7 parasites were routinely cultured in human 0+ erythrocytes at 5% hematocrit in complete RPMI 1640 medium and maintained at 37°C under 3% CO_2_, 5% O_2_. Complete RPMI medium consisted of RPMI 1640 (GlutaMAX, gibco, Thermo Fisher Scientific) supplemented with 0.5% AlboMAX II Lipid-Rich BSA (gibco, Thermo Fisher Scientific), 0.2 mM hypoxanthine (c.c. pro GmbH), 25 mM Hepes (pH 7.3; Sigma-Aldrich, Merck) and 12.5 µg/mL gentamycin (Carl Roth). Parasitemia was monitored using Giemsa-stained blood smears and parasites were maintained at parasitemias below 2%. If required, parasites were synchronized using standard sorbitol synchronization. Briefly, packed blood cells were incubated in 10 volumes of 5% sorbitol for 20 min at 37°C, then pelleted and washed once with RPMI complete before return to culture.

For transfection, 50 to 100 µg purified and ethanol-precipitated DNA were resuspended in 30 µl TE buffer and 370 µl CytoMix and mixed with 200 µl packed erythrocytes containing at least 4% ring stages. Parasites were electroporated (310 kV, 950 µF) using a Gene Pulser II (Bio-Rad) and transgenic parasites selected for by drug selection using 2.5 µM WR99210 (Jacobus Pharmaceutical Company). For genotyping transgenic parasite lines, genomic DNA was isolated from saponin-lysed and washed parasite pellets using the DNeasy Blood & Tissue Kit (Qiagen). PCR products from the genotyping reaction were sent for sequencing.

### Growth assay

To measure parasite growth, sorbitol-synchronised ring stage parasites of knockout lines and their matching controls were seeded at a parasitemia of 0.05% and 4% hematocrit into 6-well dishes. Medium was changed daily and smears and samples for flow cytometry (100 µl culture) were taken every 24 hours for three consecutive asexual cycles (6 days). To investigate formation of gametocytes, cultures were kept for additional 10 days, changing medium and collecting smears daily.

To monitor stage development, parasites were tightly synchronized to a 4 hour window. To this end, schizonts were purified by underlaying the culture with 65% Nycodenz solution (Axis-shield diagnostics) in RPMI and centrifuged for 25 min at 1000 rpm without brake. The interphase containing schizonts was collected and washed once, the parasites were mixed with medium containing blood at 4% hematocrit, transferred to a 6-well plate, and cultured for 4 h to allow for reinvasion, followed by a round of sorbitol synchronization. Parasites were followed for a full cycle, collecting blood smears and samples for flow cytometry (100 µl culture) every 8 h.

Cell samples for flow cytometry were pelleted, fixed in 200 µl 4% PFA/0.0075% Glutaraldehyde/PBS over night at 4°C and then stored in PBS. For quantification of parasitemia, cells were stained with SybrGreen (1:1000; Thermo Fisher Scientific) in PBS for 15 min at 37°C, washed once in 200 µl PBS and analysed on a BD FACS Celesta. Parasitemia was determined as percentage of SybrGreen-positive cells of all single cells. Growth rates were calculated as the average increase in parasitemia in two consecutive asexual cycles and normalized to the average growth rate of matching control parasite lines.

### Statistics

Statistical analysis was undertaken using GraphPad Prism 9.2.0 (GraphPad, San Diego, CA, USA). Calculation of the binomial confidence interval was performed using Stata/MP 17.0 for Windows (StataCorp LP, College Station, TX, USA).

