## Supplement for "Vaccination by single dose sporozoite injection of blood stage attenuated malaria parasites"

Supplementary information

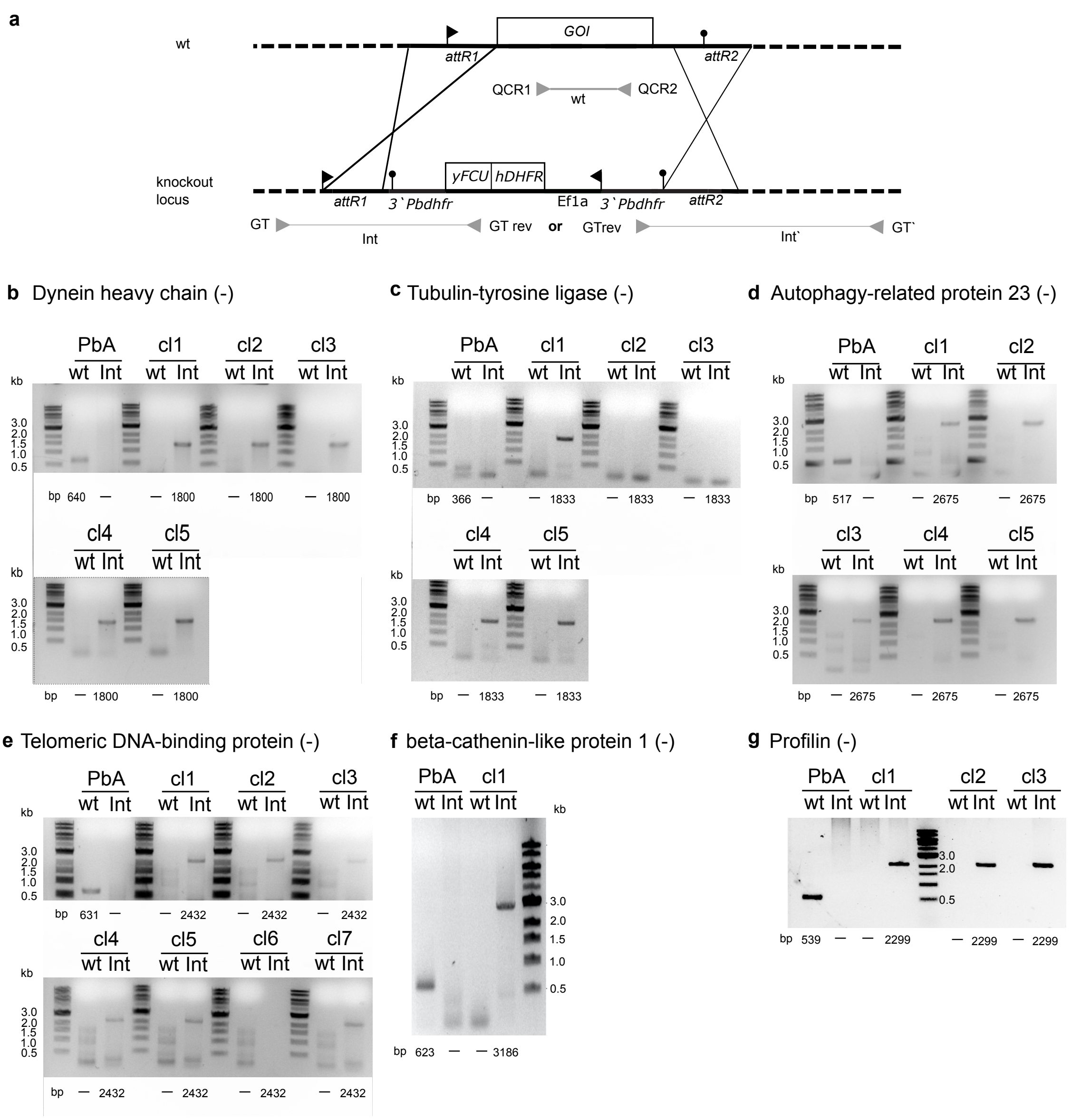


**Figure S1. Generation of *P. berghei* mutants.**
**(a)** Schematic showing chromosome locus of wildtype (top) and mutant after integration of a PlasmoGEM vector (bottom). Primers used for PCR and expected amplicons are indicated. **(b-g)** Genotyping PCRs verify correct integration for indicated clonal gene-deletion parasite lines. Expected amplicon sizes below the gels; for primer combinations see supplemental table 2. Abbreviations: GOI gene of interest; attR attachment sites for recombination; QCR quality control R; GT genotyping; rev reverse; int integration; Pbdhfr *Plasmodium berghei* dihydrofolate reductase; yFCU yeast cytosine deaminase and uridyl phosphoribosyl transferase; hDHFR human dihydrofolate reductase; cl clone; bp base pairs; kb kilo base pairs.

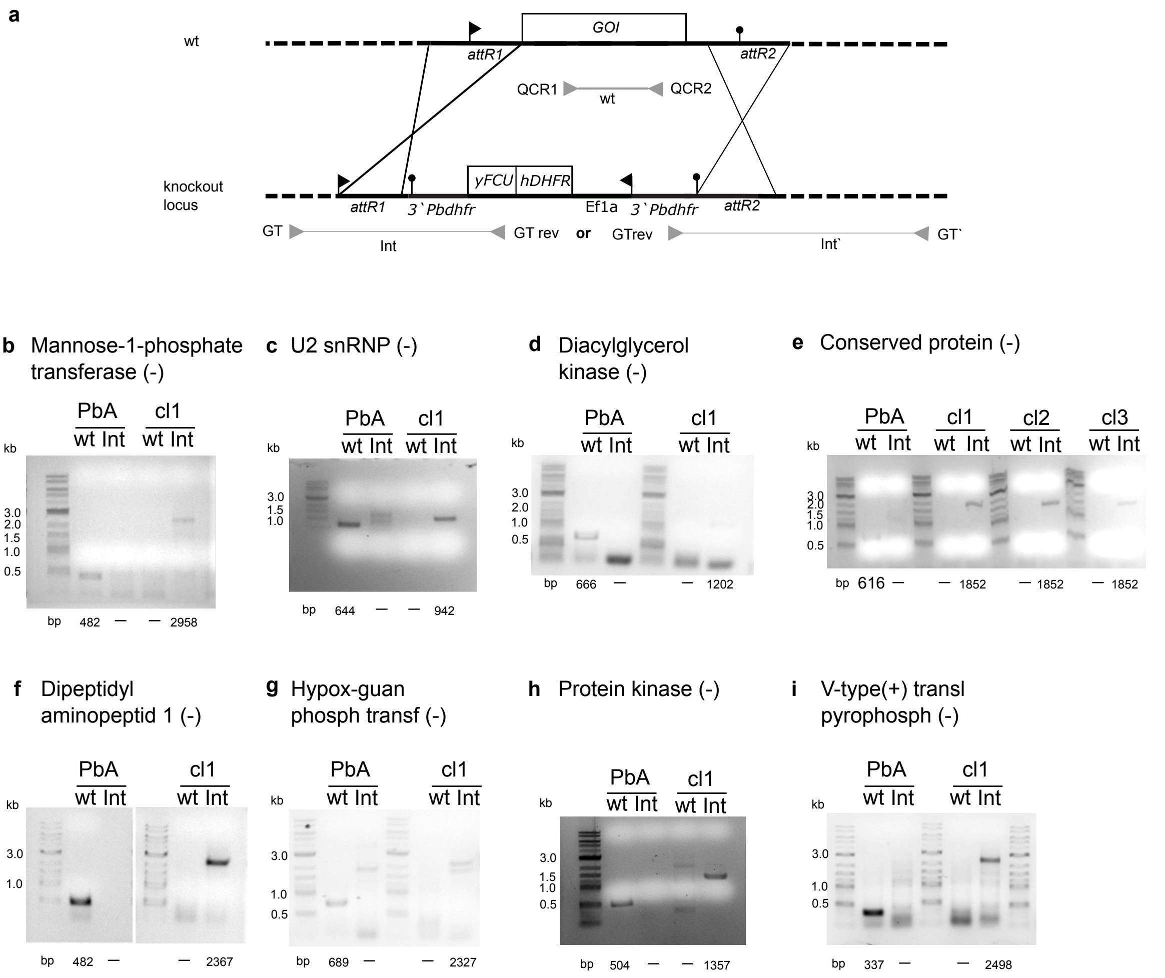


**Figure S2. Generation of *P. berghei* mutants.**
**(a)** Schematic showing chromosome locus of wildtype (top) and mutant after integration of a PlasmoGEM vector (bottom). Primers used for PCR and expected amplicons are indicated. **(b-g)** Genotyping PCRs verify correct integration for indicated clonal gene-deletion parasite lines. Expected amplicon sizes below the gels; for primer combinations see supplemental table 2. Primer combinations used can be found in supplemental table 2. Abbreviations: GOI gene of interest; attR attachment sites for recombination; QCR quality control R; GT genotyping; rev reverse; int integration; Pbdhfr *Plasmodium berghei* dihydrofolate reductase; yFCU yeast cytosine deaminase and uridyl phosphoribosyl transferase; hDHFR human dihydrofolate reductase; cl clone; bp base pairs; kb kilo base pairs.

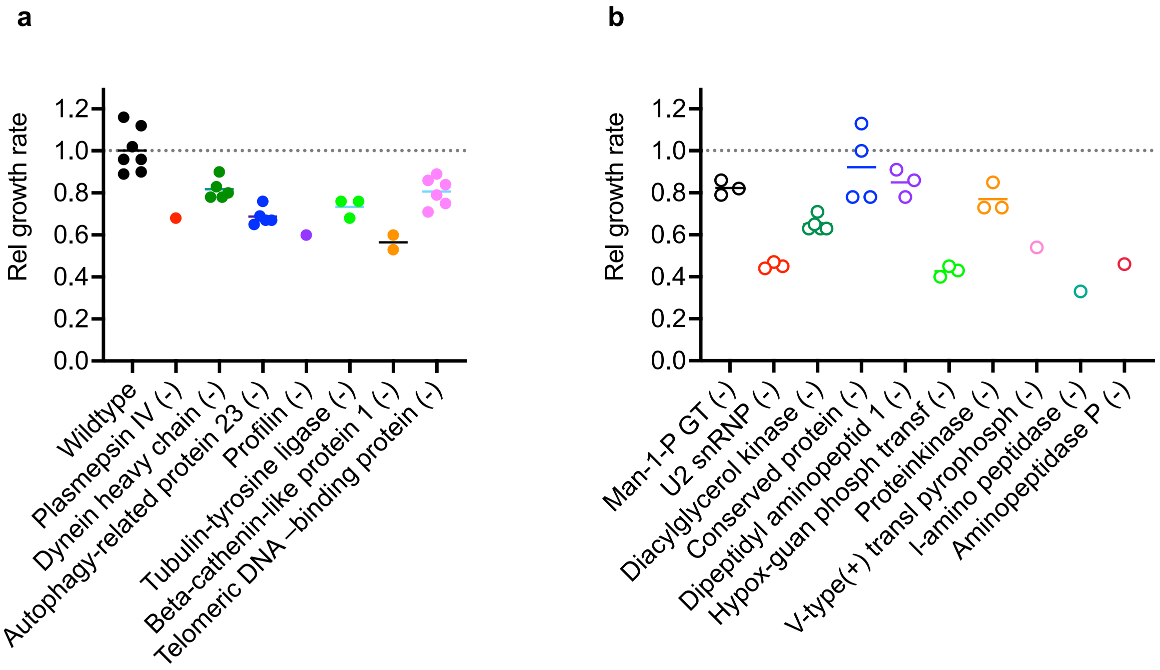


**Figure S3. Reduced relative growth rates of mutants.**
**(a, b)** Intraerythrocytic growth of mutants with a reported PlasmoGEM growth rate of 0.4-0.6 (a) and 0.2-0.4 (b). Dotted line represents average growth rate of wild type.

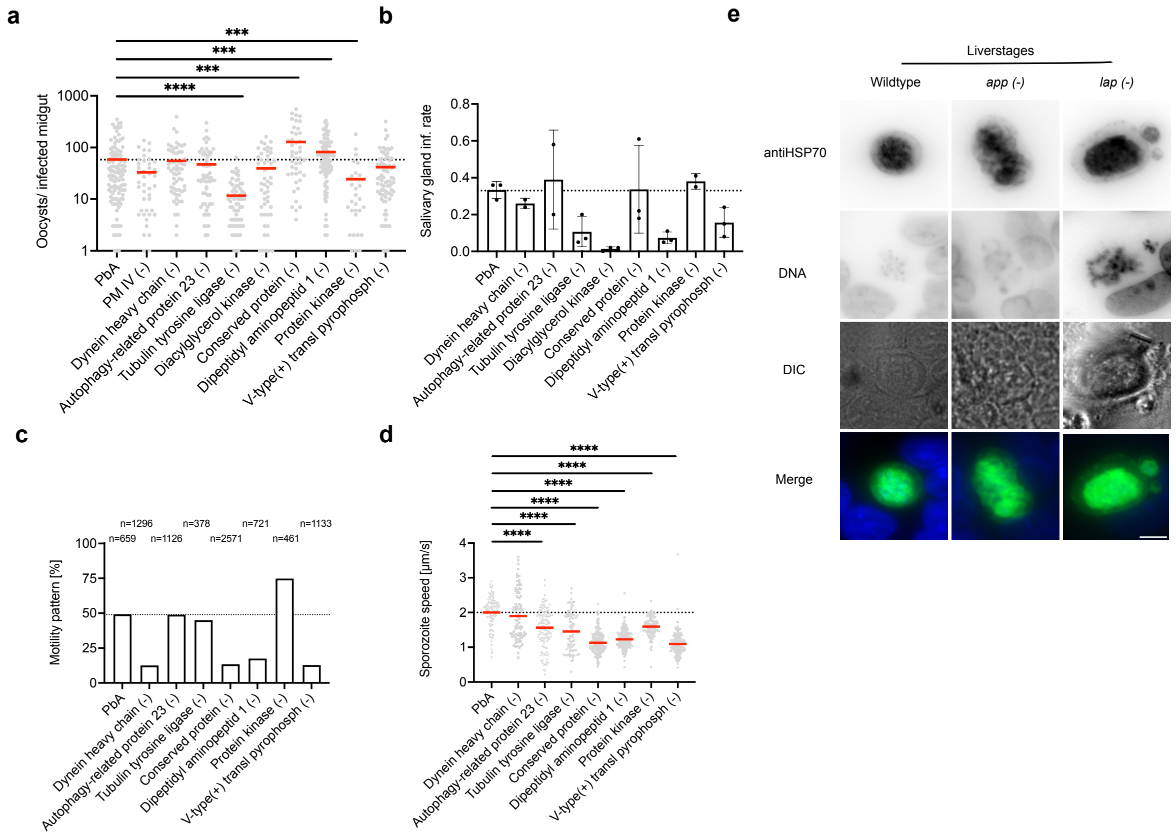


**Figure S4. Development of mutants through mosquitoes and liver stages.**
**(a)** Oocyst numbers per infected mosquito. Dotted line depicts mean oocyst numbers of wildtype parasites. Two-tailored Mann-Whitney test * indicated p< 0.05. **(b)** Salivary gland infection rate: Salivary gland derived sporozoites over midgut derived sporozoites. **(c,d)** Productive gliding motility (c) and speed (d) of salivary gland-derived sporozoites. Dotted line indicates mean of wild type sporozoites. Two-tailored Mann-Whitney test * indicated p< 0.05. **(e)** *In vitro* development of wildtype, *app(-)* and *lap(-)* liver stage parasites fixed 48h post-invasion of HepG2 cells. Nuclei were stained by Hoechst, and parasites with an anti-*Pb*HSP70 antibody. Scale bar 5 µm.

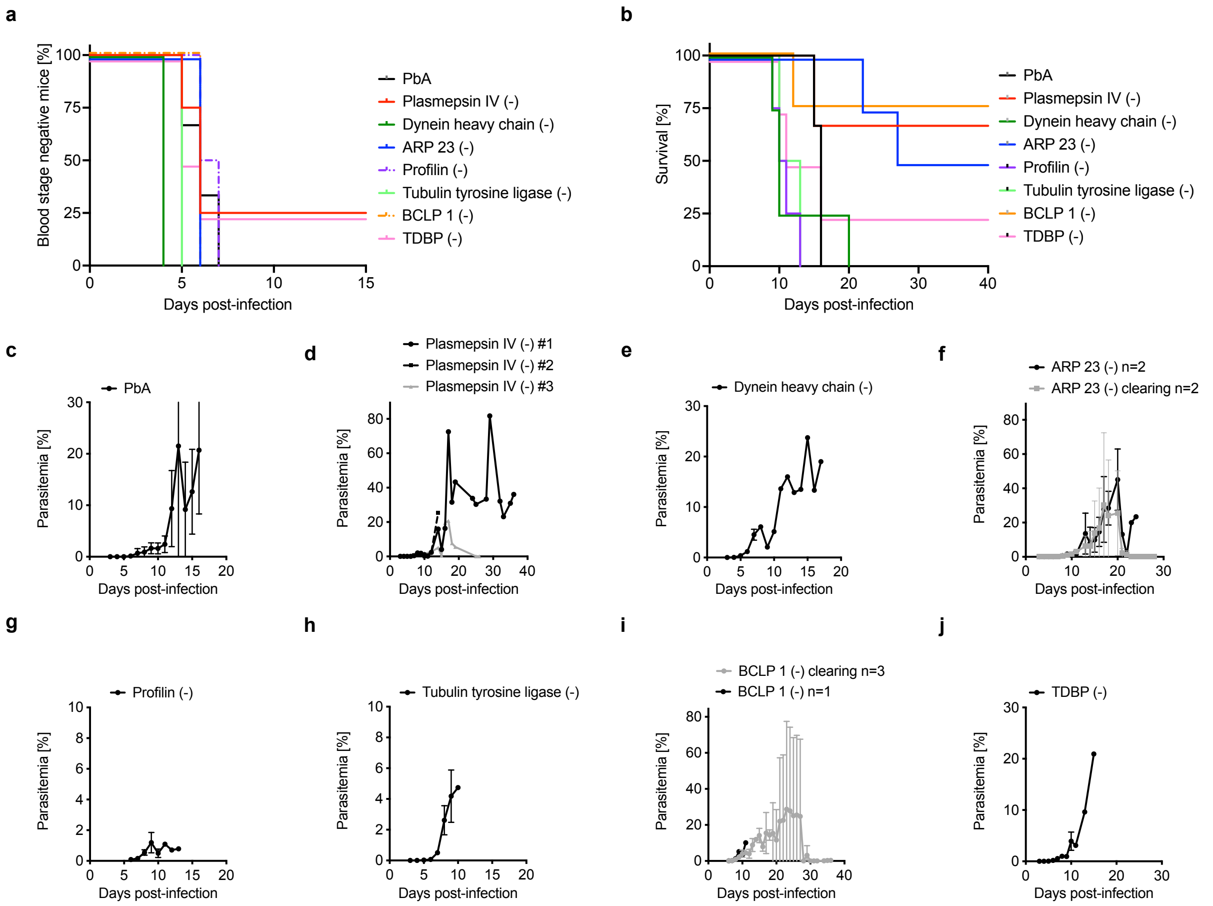


**Figure S5. Infection of Swiss mice with 100 iRBCs (mutants with an expected growth rate of 0.4-0.6).**

**(a,b)** Kaplan-Meyer plots indicating percentage of blood stage negative mice (a) and survival (b) from *i.v.* infections of 4 mice with the indicated mutants. For better visibility data were nudged to prevent overlap. **(c-j)** Course of infection with the different mutants. Mean parasitemia is shown except for (d), where individual parasitemia curves of all infected mice are displayed. ARP 23, autophagy-related protein 23; BCLP 1, beta-cathenin like protein 1; TDBP, telomeric DNA-binding protein.

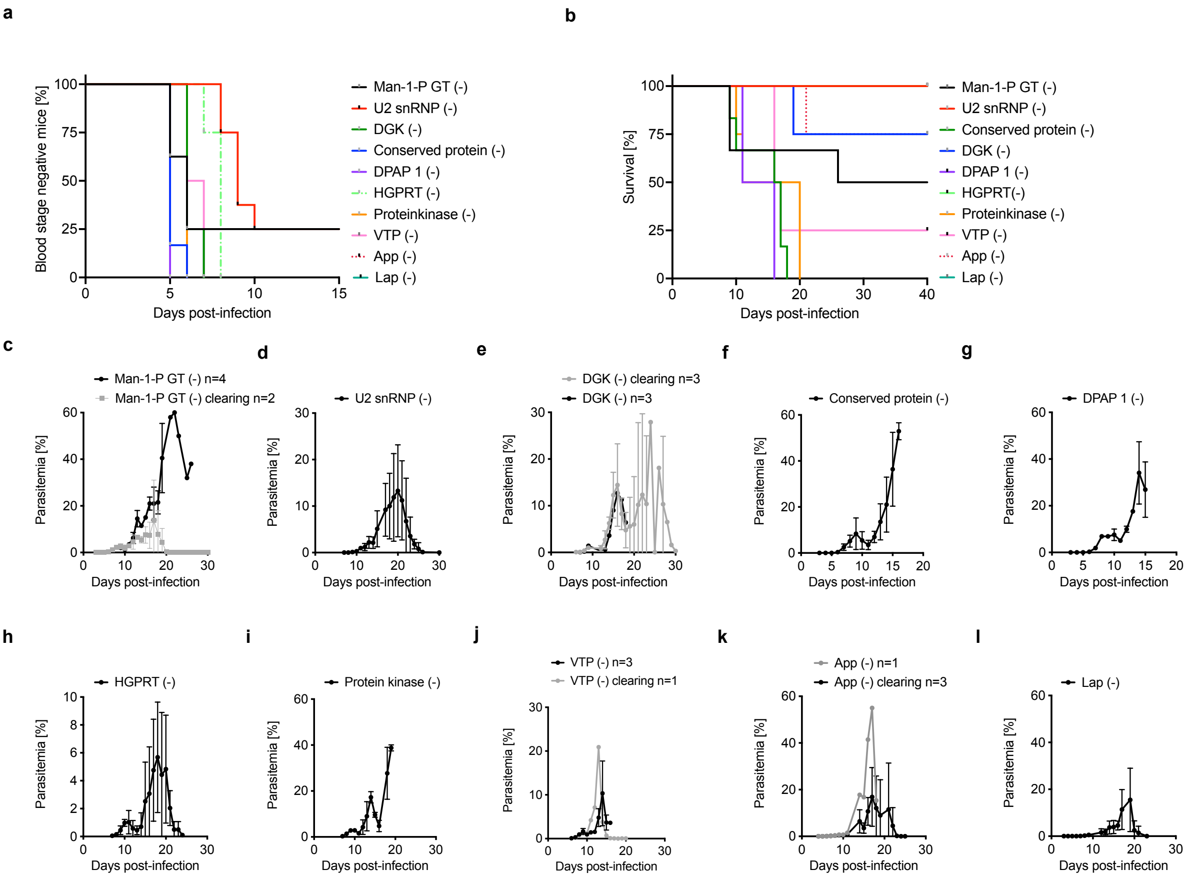


**Figure S6. Infection of Swiss mice with 100 iRBCs (mutants with an expected growth rate of 0.2-0.4).**

**(a,b)** Kaplan-Meyer plots indicating percentage of blood stage negative mice (a) and survival (b) from *i.v.* infections of 4 mice with the indicated mutants. For better visibility data were nudged to prevent overlap. **(c-j)** Course of infection with the different mutants. Mean parasitemia is shown for all mice dying (dark line) or clearing (light line) except for (d), where individual parasitemia curves of all infected mice are displayed. DGK, diacylglycerol kinase; DPAP 1, dipeptidyl aminopeptidase; HGPRT, hypoxanthine-guanidine phosphoribosyl transferase; VTP, V-type(+) pyrophosphatase.

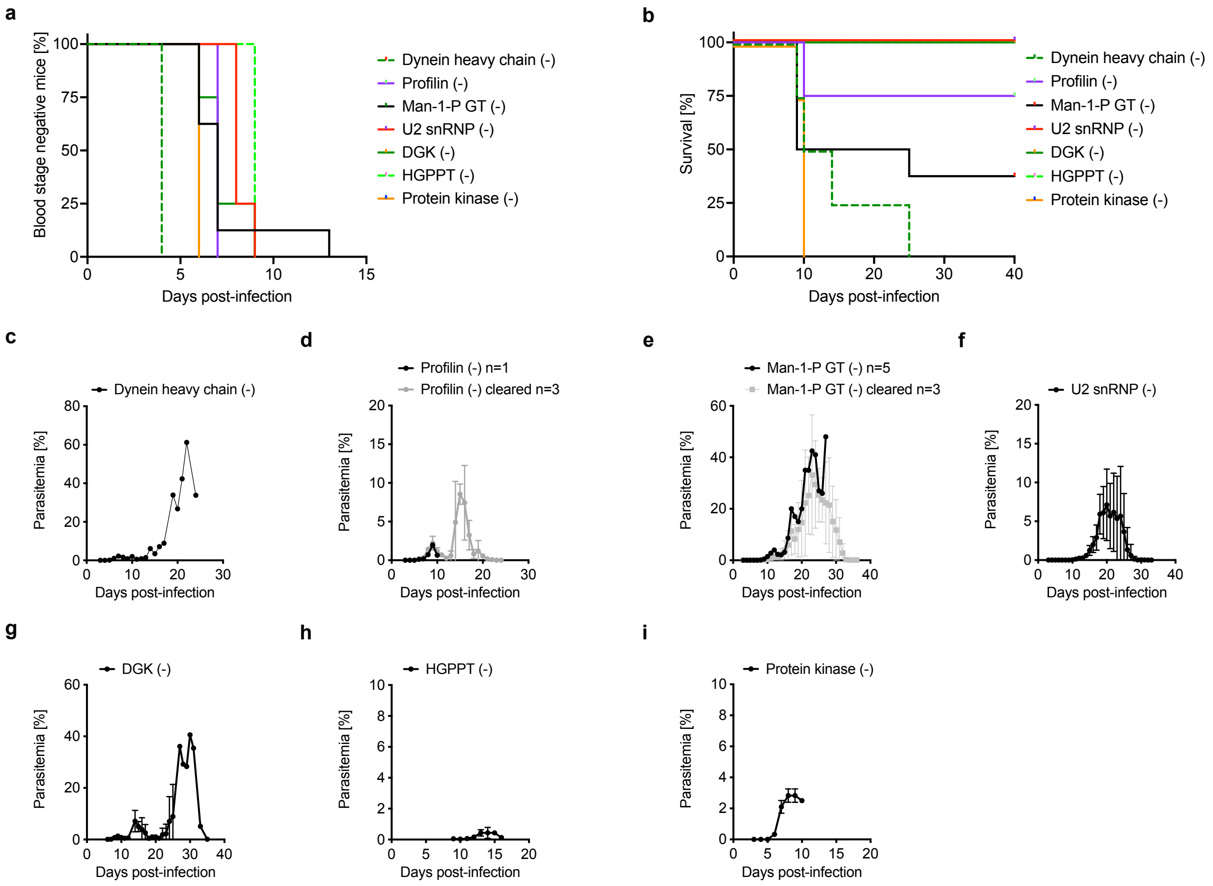


**Figure S7. Infection of C57Bl/6 mice infected with 100 iRBCs.**
**(a,b)** Kaplan-Meyer plots indicating percentage of blood stage negative mice (a) and survival (b) from i.v. infection of 4 mice for dynein heavy chain (-), profilin (-), man-1-P GT (-), DGK (-) and protein kinase (-) or 8 mice for U2 snRNP(-) and HGPRT. For better visibility data were nudged to prevent overlap. **(c-i)** Course of infection with the different mutants. Mean parasitemia is shown for of all mice dying (dark line) or clearing (light line). Note that all mice infected with *U2 snRNP (-)*, *diacylglycerol kinase (-)* or *hypox-guan phosph transf (-)* were able to survive and became blood stage negative latest by day 35. U2 snRNP**,** snRNP-associated SURP motif-containing protein; DGK, diacylglycerol kinase; HGPRT, hypoxanthine-guanidine phosphoribosyl transferase.

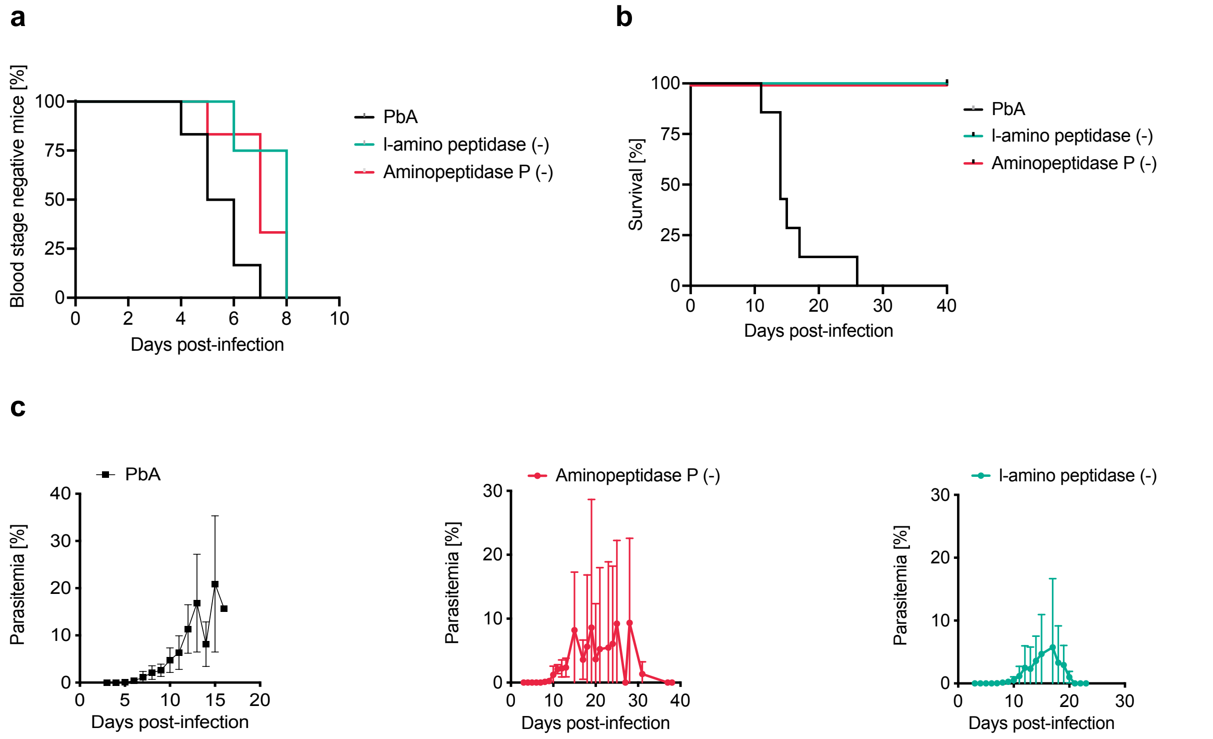


**Figure S8. Infection of Swiss mice by 10.000 salivary gland derived sporozoites. (a,b)** Kaplan-Meyer plots indicating percentage of blood stage negative mice (a) and survival (b) after i.v. infection with the indicated mutants. For better visibility data were nudged. **(c)** Parasitemia curves of infected mice are shown as mean parasitemia of all mice infected. Note that infections with *l-aminopeptidase (-)* or *aminopeptidase P (-)* were cleared and mice became blood stage negative between d25 and d39 post-infection.

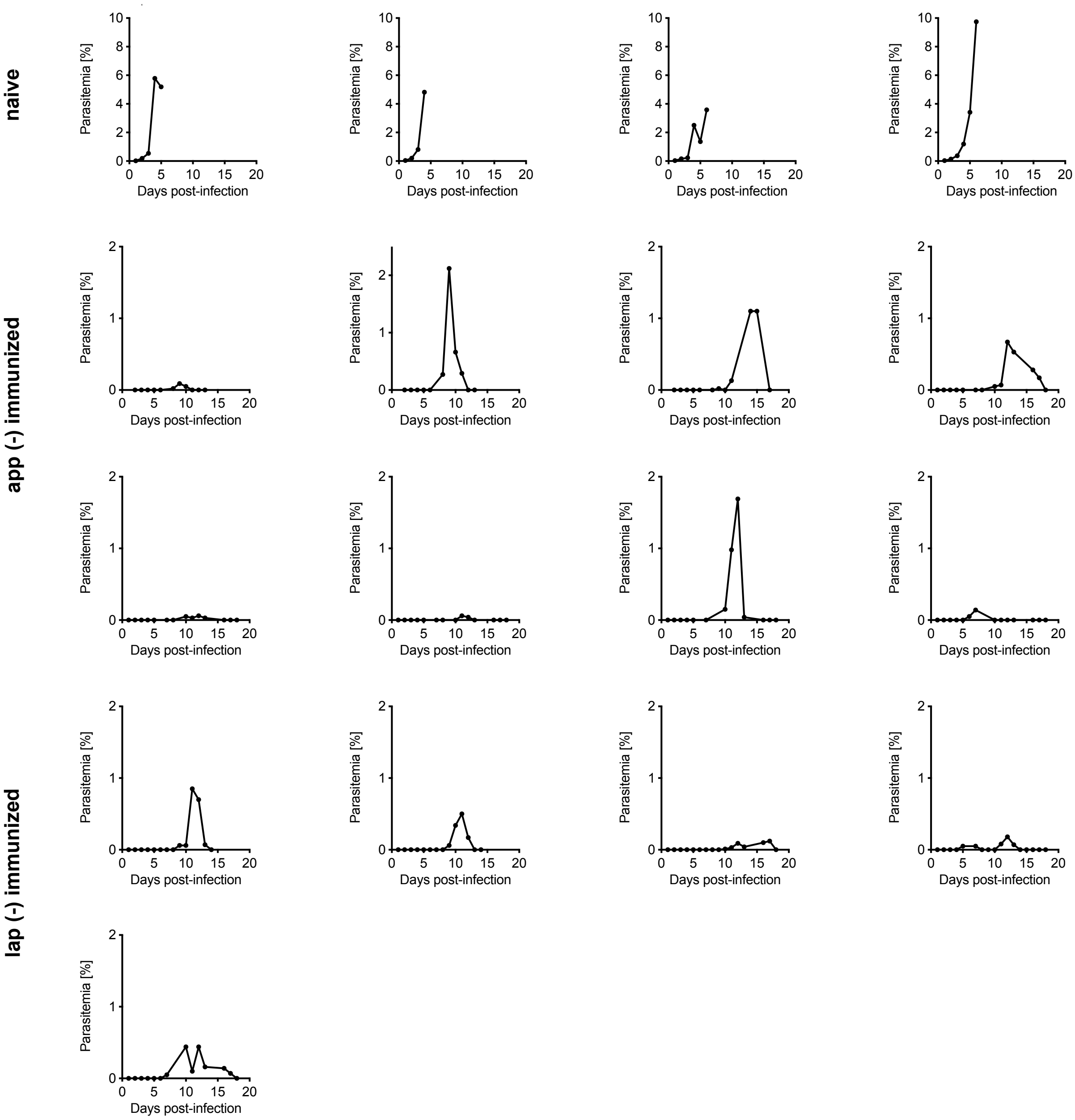


**Figure S9. Individual parasitemia curves**

Blood stage positive mice were challenged 90 days post-infection with the indicated mutants; top row: age matched naïve animals, which all died.

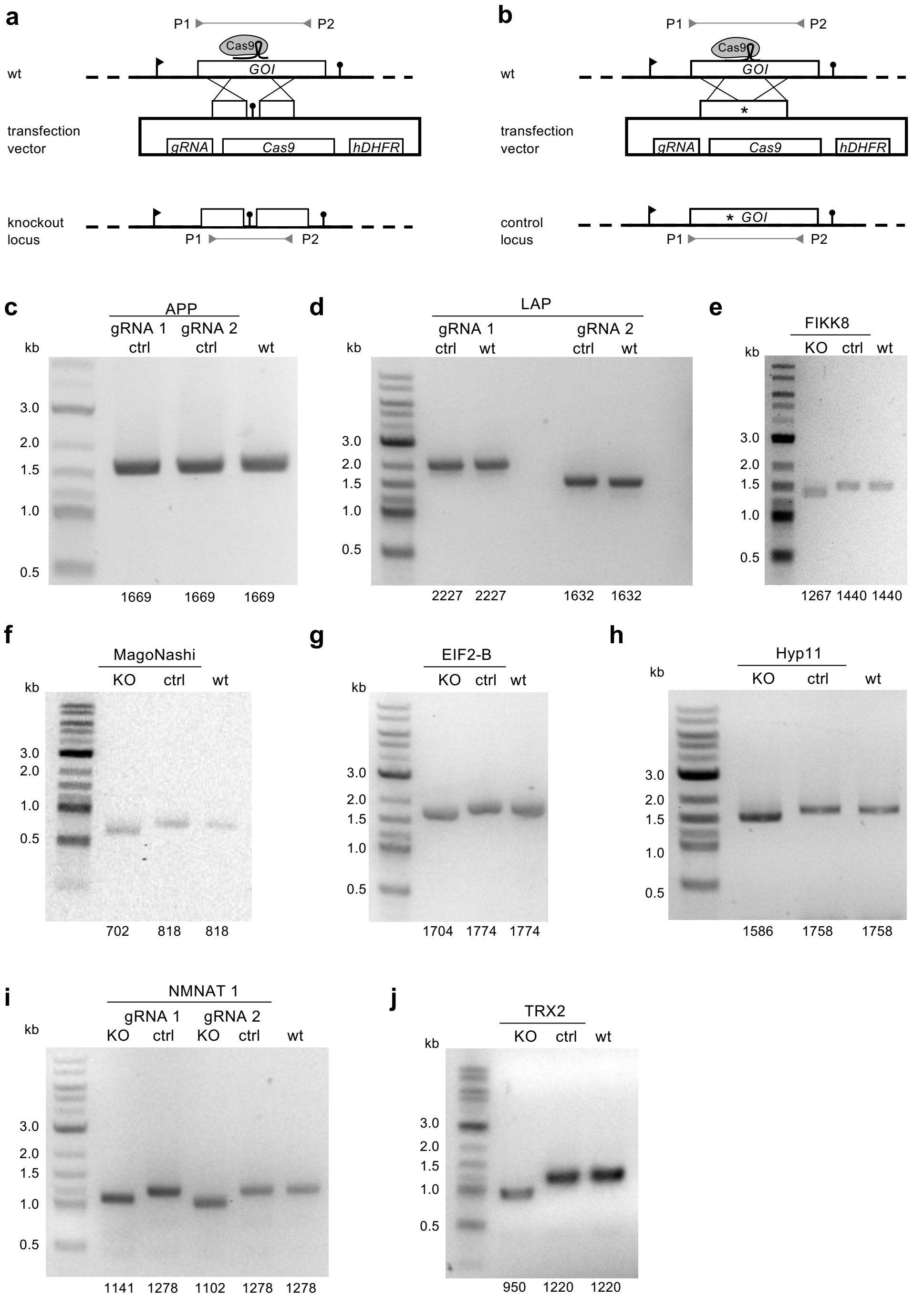


**Figure S10. Generation of slow-growing *P. falciparum* parasites.**
**(a, b)** Strategy for the generation of knockout (**a**) or control (**b**) parasite lines. The binding sites for genotyping primers P1 and P2 are indicated. GOI, gene of interest; hDHFR, human dihydrofolate reductase (resistance marker). **(c-j)** Genotyping PCRs verify the indicated knockout and control lines; expected amplicon sizes are indicated below the gels. Control PCR products were additionally sequenced to confirm introduced silent point mutations. Primer combinations: table S3. GOI, gene of interest; gRNA, guide RNA; hDHFR, human dihydrofolate reductase; KO, knockout; ctrl, control; wt, wildtype.

| Parasite line | Gene ID | | | Plasmo GEM ID |
|---|---|---|---|---|
| | P. berghei | P. falciparum | P. knowlesi | |
| Plasmepsin IV (-) | PBANKA_ 1034400 | PF3D7_ 1408100 | PKNH_ 1350300 | n.a. |
| Dynein heavy chain (-) | PBANKA_ 0615700 | PF3D7_ 0718000 | PKNH_ 0214800 | PbGEM -330283 |
| Autophagy-related protein 23 (-) | PBANKA_ 0921700 | PF3D7_ 1126700 | PKNH_ 0924600 | PbGEM -332739 |
| Profilin (-) | PBANKA_ 0833000 | PF3D7_ 0932200 | PKNH_ 0730900 | PbGEM -286426 |
| Tubulin tyrosine ligase (-) | PBANKA_ 0901900 | PF3D7_ 1147200 | PKNH_ 0945000 | PbGEM -287138 |
| Telomeric DNA – binding protein (-) | PBANKA_ 1205000 | PF3D7_ 1006800 | PKNH_ 0805600 | PbGEM -304250 |
| Beta-catenin like protein (-) | PBANKA_ 0910100 | PF3D7_ 1138600 | PKNH_ 0936800 | PbGEM -288226 |
| Man-1-P-GT (-) | PBANKA_ 1022300 | PF3D7_ 1420900 | PKNH_ 1337100 | PbGEM -037559 |
| Diacylglycerol kinase (-) | PBANKA_ 133460 | PF3D7_ 1471400 | PKNH_ 1209600 | PbGEM -265364 |
| U2 snRNP (-) | PBANKA_ 1039300 | PF3D7_ 1402700 | PKNH_ 1355400 | PbGEM -243939 |
| Conserved protein (-) | PBANKA_ 0519500 | No | No | PbGEM -533923 |
| Dipeptidyl aminopeptidase (-) | PBANKA_ 0931300 | PF3D7_ 1116700 | PKNH_ 0914400 | PbGEM -032467 |
| Hypoxanthine-guanine phosphoribosyl transferase (-) | PBANKA_ 1210800 | PF3D7_ 1012400 | PKNH_ 0812200 | PbGEM -336339 |
| Protein kinase (-) | PBANKA_ 0826900 | PF3D7_ 0926100 | PKNH_ 0724000 | PbGEM -259660 |
| V-type H(+)-translocating pyrophosphatase (-) | PBANKA_ 132050 | PF3D7_ 1235200 | PKNH_ 1454900 | PbGEM -253763 |
| Aminopeptidase P (-) | PBANKA_ 1318100 | PF3D7_ 1454400 | PKNH_ 1227700 | n.a. |
| I-aminopeptidase (-) | PBANKA_ 1309900 | PF3D7_ 1446200 | PKNH_ 1236000 | n.a. |

**Table S1. List of generated clonal *P. berghei* parasite lines**.
Gene IDs for *P. berghei* and homologues in *P. falciparum* and *P. knowlesi* as well as PlasmoGEM IDs used to generate the *Pb* lines are shown.

| Gene of interest | PlasmoGEM Vector ID | Transfection attempts | Transfection succesfull | Limiting dilutions attempts |
|---|---|---|---|---|
| Sec1 familiy protein | PbGEM-232532 | 3 | yes | 5 |
| Replication factor C subunit 2 | PbGEM-230189 | 3 | yes | 3 |
| ALP2a | PbGEM-264748 | 4 | yes | 1 |
| Calmodulin-like protein | PbGEM-330979 | 3 | yes | 4 |
| CS-domain protein | PbGEM-331875 | 1 | yes | 1 |
| calcium-dependent protein kinase 5 | PbGEM-111682 | 4 | no | n.a. |
| phosphatidylinositol N-acetylglucosaminyltransferase | PbGEM-090803 | 3 | yes | 1 |
| palmitoyltransferase DHHC4, putative | PbGEM-065186 | 2 | no | n.a. |
| Ras-related protein Rab-5B | PbGEM-109737 | 2 | no | n.a. |
| mitochondrial carrier protein, putative | PbGEM-254102 | 3 | no | n.a. |
| histone acetyltransferase | PbGEM-242177 | 2 | yes | 2 |
| coatomer subunit gamma, putative | PbGEM-260652 | 7 | no | n.a. |
| unknown | PbGEM-268387 | 4 | no | n.a. |
| tRNA-YW synthesizing protein, putative | PbGEM-261652 | 2 | no | n.a. |
| GPI-anchor transaminase | PbGEM-240110 | 3 | no | n.a. |
| WW domain-binding protein 11, putative | PbGEM-294664 | 6 | yes | 3 |
| conserved Plasmodium protein, unknown function | PbGEM-335483 | 3 | no | n.a. |
| conserved Plasmodium protein, unknown function | PbGEM-316571 | 2 | yes | 2 |
| Multidrug resistance protein | PbGEM-337155 | 2 | no | n.a. |
| cytoadherence linked asexual protein, putative | PbGEM-256118 | 3 | no | n.a. |
| conserved Plasmodium protein, unknown function | PbGEM-260988 | 3 | no | n.a. |

| | | | | |
|---|---|---|---|---|
| Sec24 subunit, putative | PbGEM-246799 | 3 | no | n.a. |
| conserved Plasmodium protein, unknown function | PbGEM-268387 | 1 | no | n.a. |
| conserved Plasmodium protein, unknown function | PbGEM-298869 | 3 | no | n.a. |
| conserved Plasmodium protein, unknown function | PbGEM-329659 | 3 | no | n.a. |
| M1-family alanyl aminopeptidase, putative | PbGEM-109859 | 1 | no | n.a. |
| mago nashi protein homologue, putative | PbGEM-235866 | 2 | no | n.a. |
| FbpA domain protein, putative | PbGEM-323051 | 1 | no | n.a. |
| U6 snRNA phosphodiesterase, putative | PbGEM-317307 | 1 | no | n.a. |
| protein kinase, putative | PbGEM-337395 | 1 | no | n.a. |
| histone acetyltransferase subunit NuA4, putative | PbGEM-321795 | 1 | no | n.a. |
| single-stranded DNA-binding protein, putative | PbGEM-246443 | 1 | no | n.a. |
| BIR protein;PIR protein | PbGEM-537051 | 1 | no | n.a. |

**Table S2. Overview of performed transfections and limiting dilutions that failed to yield clonal mutants.** No integration of the plasmid could be verified after transfection or the limiting dilution failed to yield a clonal population. Abbreviations: n.a. not applicable.

| Gene ID | Primer | Sequence (5‘ → 3‘) |
|---|---|---|
| PBANKA_0615700 (Dynein heavy chain ) | GT | ACATTTCACGATGGCGCGGA |
| | GT rev | CTTTGGTGACAGATACTAC |
| | QCR1 | TCCCCTTTGTGCCTTCAGAGCT |
| | QCR2 | AGGAATCACCACGAAAGGGACA |
| PBANKA_0921700 (Autophagy-related protein 23 ) | GT | ACGTGATGTGAATGCCTACA |
| | GT rev | TGATTAGCATAGTTAAATAAAAAAAGTTG |
| | QCR1 | TCTTCACATTCTTCTTGCATGT |
| | QCR2 | TTCACCCCCTACACCGATAT |
| PBANKA_0833000 (Profilin ) | GT | TGGCACACTTGGTTTGACAGAGGT |
| | GT rev | CATACTAGCCATTTTATGTG |
| | QCR1 | TCATCGGGGGTTGCTACGCA |
| | QCR2 | GCGTAAGGCTTCGTCCGTTT |
| PBANKA_0901900 (Tubulin tyrosine ligase ) | GT | GGTTGTGCAAACCGAGACTT |
| | GT rev | TGATTAGCATAGTTAAATAAAAAAAGTTG |
| | QCR1 | TTGGGTTGAGCCAGATTCGA |
| | QCR2 | AGCATATTAGAGTCACAACCA |
| PBANKA_1205000 (Telomeric DNA-binding protein ) | GT | TGCAGTCGTATTTGTCACCACA |
| | GT rev | TGATTAGCATAGTTAAATAAAAAAAGTTG |
| | QCR1 | ACGTCATTACATTTGTGGTAGCCA |
| | QCR2 | ACCACTGAAGAAGGCGAAGA |
| PBANKA_0910100 (Beta-catenin like Protein 1) | GT | TGATGTGATGAGCAAATCCGA |
| | GT rev | TGATTAGCATAGTTAAATAAAAAAAGTTG |
| | QCR1 | CGTCAATGTCAATCTCATCTTCATCGC |
| | QCR2 | TCTGTTCCAGAAAATGGGGT |

| | | |
|---|---|---|
| PBANKA_1022300 (Man-1-P-GT ) | GT | ACTTTGATGCCTGGCTCTCCT |
| | GT rev | CTTTGGTGACAGATACTAC |
| | QCR1 | TGGGCAGATATTGGAAAACCTTCTGA |
| | QCR2 | TCAATTCGAGCCCAGCTCCCT |
| PBANKA_1333460 Diacylglycerol kinase | GT | ACCAGAGGGTTGCCATTTGCACA |
| | GT rev | CTTTGGTGACAGATACTAC |
| | QCR1 | AGCCCCAAATCAACACCTGAACGA |
| | QCR2 | TGGGTGGAAGAAACCAATTGGGA |
| PBANKA_0519500 (U2 snRNP ) | GT | ACCCTCGTTGCTCACATAACCGA |
| | GT rev | CATACTAGCCATTTTATGTG |
| | QCR1 | CGGCATCTCCCTCAAACGACCG |
| | QCR2 | ACGAAAACGCCCTTCACATCT |
| PBANKA_0519500 (Conserved protein ) | GT | TTGGAGGCGCTCTCATTGGT |
| | GT rev | GGCTATTCATACTAGCCATTTTATGTGTG |
| | QCR1 | TGTGCTTTGACGGTTTAGCTCC |
| | QCR2 | AAGAGACGCAAAAAGCACAC |
| PBANKA_093130 0 (Dipeptidyl aminopeptidase) | GT | GCTGCTATCAATGCAGCACCACCA |
| | GT rev | CTTTGGTGACAGATACTAC |
| | QCR1 | ACACGTGGTGCTGGTAAAGTGCT |
| | QCR2 | TTAATGCAAGGGAGCCGACC |
| PBANKA_121080 0 (Hypoxanthine-guanine | GT | CCATCTTTCAAATGGCCCTCG |
| | GT rev | CATACTAGCCATTTTATGTG |
| | QCR1 | AAAAGGGGCGTGCACAAATA |
| | QCR2 | TGTTGCATGCGTTCCAGTTGT |

| | | |
|---|---|---|
| PBANKA_0826900 (Protein kinase ) | GT | TGGGAATTGCGCATTTGGTGATCG |
| | GT rev | CATACTAGCCATTTTATGTG |
| | QCR1 | TGTACCATCTTCCTCTGATT |
| | QCR2 | GGAGGAATAGCCACATTACCTTCACG |
| PBANKA_132050 (V-type H(+)-translocating pyrophosphatase) | GT | AGTCATTTAGGAGCAGCGGA |
| | GT rev | CTTTGGTGACAGATACTAC |
| | QCR1 | CGAGAGCGGGAGGACCAAAT |
| | QCR2 | ATTGCCGTTCTAAAGCACTT |
| PF3D7_1454400 (APP) | gRNA1 F | TATTTACCGTTGATGTTAACATGA |
| | gRNA1 R | AAACTCATGTTAACATCAACGGTA |
| | gRNA2 F | TATTATCCATCATGTTAACATCAA |
| | gRNA2 R | AAACTTGATGTTAACATGATGGAT |
| | HR1 For | GAGGTACCGAGCTCGAATTCAGAGGCAAAATTTAGTATCGG |
| | HR1 KO Rev | CACTATTAATTAAATTATTATTAATATATATTCTTAAATATATTGGAAC CAC |
| | HR1 Ctrl gRNA1 Rev | GATTATTATCCATCATATTCACATCAACGGTATTATTATTACTCATC |
| | HR1 Ctrl gRNA1 Rev | CCATCATGTTAACATCAACTGTATTATTATTACTCATCATTTCGG |
| | HR2 KO For | TAAGAATATATATTAATAATAATTTAATTAATAGTGATGAACACAATTC TG |
| | HR2 Ctrl gRNA1 For | ATAATACCGTTGATGTGAATATGATGGATAATAATCCTGCTGC |
| | HR2 Ctrl gRNA2 For | TGATGAGTAATAATAATACAGTTGATGTTAACATGATGGATAATAATC |
| | HR2 Rev | CGAAAAGTGCCACCTGACGTCGGTGAATATTGATAATCATAACCTC |
| | Genotyping P1 For | GGTAAGAGGCAAAATTTAGTATCGG |
| | Genotyping P2 Rev | TAGTTCCATGTAAATATTGTCCTCCC |
| PF3D7_14462 00 (LAP) | gRNA1 F | TATTGCTGTTGGTTATGTAGGATG |
| | gRNA1 R | AAACCATCCTACATAACCAACAGC |
| | gRNA2 F | TATTGGAGGTTGTAATGTTGAAGA |
| | gRNA2 R | AAACTCTTCAACATTACAACCTCC |

| | | |
|---|---|---|
| | HR1 gRNA1 For | GAGGTACCGAGCTCGAATTCTCCTAAAATCTACTATACACAGTGG |
| | HR1 KO gRNA1 Rev | ATCAGCAACTGATCTTATTAAACTGGCCCATTTTCTTTACCAG |
| | HR1 Ctrl gRNA1 Rev | CAACTGATCCACACCCGACGTACCCAACAGCAACTGAATTTTTATTATC |
| | HR2 KO gRNA1 For | AAAATGGGCCAGTTTAATAAGATCAGTTGCTGATTTAAGTGAAGC |
| | HR2 Ctrl gRNA1 For | TTGCTGTTGGGTACGTCGGGTGTGGATCAGTTGCTGATTTAAGTG |
| | HR2 gRNA1 Rev | CGAAAAGTGCCACCTGACGTCGCCACAATAGATGAAGCTTTAACAC |
| | HR1 gRNA2 For | GAGGTACCGAGCTCGAATTCGCCTTTGTGCATGGTTAATAATAATG |
| | HR1 KO gRNA2 Rev | ATTTTCTTTACCAGTTATTATGGATCTAAGGAAACAACTTGTGG |
| | HR1 Ctrl gRNA2 Rev | TTAATCCTTCCTCGACGTTGCAACCTCCTTTTATATCATAAACCTGAAC |
| | HR2 KO gRNA2 For | TTTCCTTAGATCCATAATAACTGGTAAAGAAAATGGGCCAG |
| | HR2 Ctrl gRNA2 For | AAGGAGGTTGCAACGTCGAGGAAGGATTAACTATTTTCTTAGTTAATAATCCTGG |
| | HR2 gRNA2 Rev | CGAAAAGTGCCACCTGACGTCCCCACAGATAAATAGGCTCCC |
| | Genotyping P1 For | ATCCTAAAATCTACTATACACAGTGG |
| | Genotyping P2 Rev | CAATATCAATATGTGCCCAAGC |
| PF3D7_0805700 (FIKK8) | gRNA F | TATTATGGATCCTAAAGATTGTTC |
| | gRNA R | AAACGAACAATCTTTAGGATCCAT |
| | HR1 For | GAGGTACCGAGCTCGAATTCAAGTAATGTTGATATGTGTGAGGG |
| | HR1 KO Rev | ATCTTCATCCTGTTCCTTATTATTAATCCTTATTTGTATCTTCCTTCATC |
| | HR1 Ctrl Rev | TCATTAAAATTTATTCCGCTACAATCTTTAGGATCCATAGAAATATTTC |
| | HR2 KO For | AGATACAAATAAGGATTAATAATAAGGAACAGGATGAAGATGAAGG |
| | HR2 Ctrl For | TGGATCCTAAAGATTGTAGCGGAATAAATTTTAATGAAATG |
| | HR2 Rev | CGAAAAGTGCCACCTGACGTCCCTTAACATCCACTTGTTCATAAC |
| | Genotyping P1 For | ATGAATGATGCTGAAGATGTGTC |

| | | |
|---|---|---|
| | Genotyping P2 Rev | GCATATCATTTTGACCTGCTCC |
| PF3D7_0725200 (Mago Nashi) | gRNA F | TATTACACTGTTTAACATCTGATA |
| | gRNA R | AAACTATCAGATGTTAAACAGTGT |
| | HR1 For | GAGGTACCGAGCTCGAATTCCCTTTAGATATTAGTAATTGAAATAAGAGC |
| | HR1 KO Rev | TCTGGGTCATTATTATTACACTTTACATATTTCAGATTCTTCAATTATACG |
| | HR1 Ctrl Rev | CTTGTTTTCCAATCTTATCAGGCATAGGCCATTCTTTATCAC |
| | HR2 KO For | TATGTAAAGTGTAATAATAATGACCCAGAAGGTTTACGTG |
| | HR2 Ctrl For | CTACTTCTAAAATTGGATCGTTATCAGATGTTAAACAGTGTAGTG |
| | HR2 Rev | CGAAAAGTGCCACCTGACGTCTTAATAAGGTACATTTACGTTCTTGG |
| | Genotyping P1 For | AATGTCTAAGAAAGATAAATTTTCCTTTAG |
| | Genotyping P2 Rev | TTAATAAGGTACATTTACGTTCTTGG |
| PF3D7_1148800 (Hyp11) | gRNA F | TATTAAGGTATAGAAGAATAGTAG |
| | gRNA R | AAACCTACTATTCTTCTATACCTT |
| | HR1 For | GAGGTACCGAGCTCGAATTCTGTGGTTGCTTGATCTGGAC |
| | HR1 KO Rev | TTGCATATTTACCTTTATTATGTTTCATTGAAAGGGTAATATAGTTG |
| | HR1 Ctrl Rev | CATGTTGTTCTTCGGCGACGATTCTTCTATACCTTGAGCATGTTTC |
| | HR2 KO For | CTTTCAATGAAACATAATAAAGGTAAATATGCAACAAGATGGTG |
| | HR2 Ctrl For | GGTATAGAAGAATCGTCGCCGAAGAACAACATGAAATTGAATTAGACG |
| | HR2 Rev | CGAAAAGTGCCACCTGACGTCCCATTAAGGTAATAATTTACTTCCTCCATC |
| | Genotyping P1 For | TGTGGTTGCTTGATCTGGAC |
| | Genotyping P2 Rev | TCTTTCTAAATAATGTTCCGGTTCATC |
| PF3D7_0828500 (EIF-2B) | gRNA F | TATTTTACTGGAGTCGCTAAAATG |
| | gRNA R | AAACCATTTTAGCGACTCCAGTAA |
| | HR1 For | GAGGTACCGAGCTCGAATTCCCTATTCAACCTTTGCCAAGAG |
| | HR1 KO Rev | ATCTTCTGTGGTGCTTATTAGCAACTGTTCAATGTCTTTGC |

| | | |
|---|---|---|
| | HR1 Ctrl Rev | AATTACTGGAGTCTGAGAAGTGAGGACATTCATTTAATAAGCAACTG |
| | HR2 KO For | CATTGAACAGTTGCTAATAAGCACCACAGAAGATCACAATG |
| | HR2 Ctrl For | ATGAATGTCCTCACTTCTCAGACTCCAGTAATTTGGAATACCG |
| | HR2 Rev | CGAAAAGTGCCACCTGACGTCAAATGATACTAGTACCTACATGCCC |
| | Genotyping P1 For | CCTATTCAACCTTTGCCAAGAG |
| | Genotyping P2 Rev | TTCGAATGATGATGATAATGATGC |
| PF3D7_1327600 (NMNAT) | gRNA1 F | TATTATTTCGTGAGCATATGTAAT |
| | gRNA1 R | AAACATTACATATGCTCACGAAAT |
| | gRNA2 F | TATTTCTGTGATGAAATTCTGTTA |
| | gRNA2 R | AAACTAACAGAATTTCATCACAGA |
| | HR1 gRNA1 For | GAGGTACCGAGCTCGAATTCGACCCTTTATCCTTCCTATAACCC |
| | HR1 KO gRNA1 Rev | TATGTCTGTGATGATTATTACTTATGCATTTTAAAGGGACCAC |
| | HR1 Ctrl gRNA1 Rev | CCATTTCGTGAGCGTACGTGATTGGATCAAAGGACCCTCC |
| | HR2 KO gRNA1 For | TTAAAATGCATAAGTAATAATCATCACAGACATAACATGTTTACC |
| | HR2 Ctrl gRNA1 For | CCTTTGATCCAATCACGTACGCTCACGAAATGGTTCTTG |
| | HR2 gRNA1 Rev | CGAAAAGTGCCACCTGACGTCTGCATAATACAAGTACGAGAAGGG |
| | HR1 gRNA2 For | GAGGTACCGAGCTCGAATTCGACCCTTTATCCTTCCTATAACCC |
| | HR1 KO gRNA2 Rev | TATGAGATTCAAGATTATTATGTAATTGGATCAAAGGACCCTC |
| | HR1 Ctrl gRNA2 Rev | TGATGAAATTCTGTCAATGATTTATCATTTCTACATCTACATATAACTA CCC |
| | HR2 KO gRNA2 For | TTGATCCAATTACATAATAATCTTGAATCTCATAGTGAAATGACTCC |
| | HR2 Ctrl gRNA2 For | GTAGAAATGATAAATCATTGACAGAATTTCATCACAGACATAACATG |
| | HR2 gRNA2 Rev | CGAAAAGTGCCACCTGACGTCTGCATAATACAAGTACGAGAAGGG |
| | Genotyping P1 For | GACCCTTTATCCTTCCTATAACCC |

| | | |
|---|---|---|
| | Genotyping P2 Rev | TGTATATATGTGCAATGGGTTAATGTTC |
| PF3D7_1345100 (TRX2) | gRNA F | TATTAAAGCGCATATACTTATTAA |
| | gRNA R | AAACTTAATAAGTATATGCGCTTT |
| | HR1 For | GAGGTACCGAGCTCGAATTCGTGTACGATGTTACATGTACAAAAG |
| | HR1 KO Rev | CATTGCTTGAGACATTATTAACAAGCTTGACACCTACACAC |
| | HR1 Ctrl Rev | TTGTCTAGGTCAACTTTGAGTAAGTATATGCGCTTTCCGTAATATTTTTG |
| | HR2 KO For | GGTGTCAAGCTTGTTAATAATGTCTCAAGCAATGATTTGATAGC |
| | HR2 Ctrl For | AGCGCATATACTTACTCAAAGTTGACCTAGACAAAAATGAATCAC |
| | HR2 Rev | CGAAAAGTGCCACCTGACGTCATATAGACCTGTTGCATTTGCC |
| | Genotyping P1 For | CTATAGCATATATAAAGATTATATATCGAAACAAC |
| | Genotyping P2 Rev | ATATAGACCTGTTGCATTTGCC |

**Table S3. Primers used in this study.**

| | PlasmoGEM ID | Design warning | Coding sequence remaining in genome (%) |
|---|---|---|---|
| Dynein heavy chain (-) | PbGEM-330283 | Yes | 25 |
| Autophagy-related protein 23 (-) | PbGEM-332739 | yes | 15 |
| Profilin (-) | PbGEM-286426 | no | n.a. |
| Tubulin tyrosine ligase (-) | PbGEM-287138 | yes | 20 |
| Telomeric DNA-binding protein (-) | PbGEM-304250 | no | n.a. |
| Beta-catenin like Protein 1 (-) | PbGEM-288226 | no | n.a. |
| Man-1-P-GT (-) | PbGEM-037559 | yes | 48 |
| Diacylglycerol kinase (-) | PbGEM-265364 | yes | 56 |
| U2 snRNP (-) | PbGEM-243939 | yes | 33 |
| Conserved protein (-) | PbGEM-533923 | no | n.a. |
| Dipeptidyl aminopeptidase (-) | PbGEM-032467 | no | n.a. |
| Hypoxanthine-guanine phosphoribosyl Transferase (-) | PbGEM-336339 | yes | <1 |
| Protein kinase (-) | PbGEM-259660 | yes | 21 |
| V-type H(+)-translocating pyrophosphatase (-) | PbGEM-253763 | no | n.a. |

**Table S4. Overview of design warnings for all generated mutants.**

| | *In vivo* multiplication rate | | | | Oocysts/ infected midgut (mean) | Infected midguts/ midguts analyzed [%] | salivary gland sporozoites / mosquito |
|---|---|---|---|---|---|---|---|
| | Plasmo GEM | CI | clone | CI | | | |
| PbA | 1 | n.a. | 1±0.11 | 0.79-1.21 | 58 | 88 | 6.500 – 27.500 |
| Plasmepsin IV (-) | 0.67 | 0.66 – 0.69 | 0.58# | n.a. | 33 | 57 | 0 |
| Dynein heavy chain (-) | 0.6 | 0.22 – 0.97 | 0.8±0.05 | 0.72-0.91 | 55 | 68 | 11.000 – 23.000 |
| Autophagy-related protein 23 (-) | 0.58 | 0.49 – 0.68 | 0.7±0.04 | 0.61-0.77 | 47 | 82 | 14.000 – 15.000 |
| Profilin (-) | 0.57 | 0.45 – 0.69 | 0.6 | n.a. | 15 | 35 | 0 |
| Tubulin tyrosine ligase (-) | 0.51 | -0.23 – 1.26 | 0.7±0.04 | 0.65-0.82 | 12 | 65 | 200 – 6.000 |
| Telomeric DNA – binding protein (-) | 0.44 | 0.33 – 0.56 | 0.8±0.07 | 0.67-0.94 | 0 | 0 | 0 |
| Beta-catenin like protein 1 (-) | 0.42 | 0.35 – 0.49 | 0.6±0.05 | 0.46-0.67 | 0 | 0 | 0 |
| Man-1-P-GT (-) | 0.39 | 0.27 – 0.52 | 0.8±0.04 | 0.85-0.89 | 0 | 0 | 0 |
| Diacylglycerol kinase (-) | 0.37 | 0.25 – 0.48 | 0.7±0.05 | 0.56-0.75 | 39 | 72 | 0 – 900 |
| U2 snRNP (-) | 0.39 | 0.31 – 0.48 | 0.5±0.02 | 0.42-0.48 | 0 | 0 | 0 |
| Conserved protein (-) | 0.35 | -0.12 – 0.82 | 0.9±0.18 | 0.58-1.27 | 119 | 76 | 5.500 – 15.000 |
| Dipeptidyl aminopeptidase (-) | 0.33 | -0.12 – 0.78 | 0.9±0.07 | 0.72-0.98 | 82 | 76 | 850 – 7.000 |
| Hypoxanthine-guanine phosphoribosyl transferase (-) | 0.29 | 0.12 – 0.45 | 0.4±0.03 | 0.38-0.48 | 0 | 0 | 0 |
| Protein kinase (-) | 0.29 | 0 – 0.58 | 0.8±0.07 | 0.63-0.91 | 24 | 70 | 5.500 – 11.000 |
| V-type H(+)-phosphoribosyl transferase (-) | 0.27 | 0.22 – 0.32 | 0.5 | n.a. | 42 | 72 | 550 – 5.100 |
| Aminopeptidase P (-) | 0.21 | 0.07 – 0.35 | 0.46* | n.a. | 25 | 65 | 1.600 – 11.000 |
| l-aminopeptidase (-) | 0.25 | 0.14 – 0.36 | 0.33* | n.a. | 26 | 52 | 1.500 – 6.000 |

**Table S5. Infection overview across life cycle**

For in vivo multiplication rates, growth rates from limiting dilutions were normalized to intra experimentally obtained wildtype multiplication rates. # indicate multiplication rate as reported in Spaccapelo et al 2010 normalized to reported wildtype multiplication rate. * indicate multiplication rates as reported in Zheng et al 2015 normalized to reported wildtype multiplication rate. Mean sporozoite numbers from at least two independent mosquito infections. Abbreviations: CI for confidence interval, n.d. not determined.

| | 100 iRBC Swiss | | | 100 iRBC C57Bl/6 | | |
|---|---|---|---|---|---|---|
| | Mice clearing/ mice infected (prepatency d) | Peak parasitemia cleared [%] (mean d) | Blood stage cleared (mean d) | Mice clearing/ mice infected (prepatency d) | Peak parasitemia cleared [%] (mean d) | Blood stage cleared (mean d) |
| Plasmepsin IV (-) | 1/3 (6) | 21 (17) | 25 | n.d. | n.a. | n.a. |
| Dynein heavy chain (-) | 0/4 (4) | n.a. | n.a. | n.d. | n.a. | n.a. |
| Autophagy-related protein 23 (-) | 2/4 (6) | 32 (19) | 23 | n.d. | n.a. | n.a. |
| Profilin (-) | 0/4 (7) | n.a. | n.a. | 3/4 (6) | 10 (15) | 23 |
| Tubulin tyrosine ligase (-) | 0/4 (5) | n.a. | n.a. | n.d. | n.a. | n.a. |
| Telomeric DNA-binding protein (-) | 0/3 (5) | n.a. | n.a. | n.d. | n.a. | n.a. |
| Beta-catenin like Protein 1(-) | 3/4 (6) | 42 (19) | 37 | n.d. | n.a. | n.a. |
| Man-1-P-GT (-) | 0/3 (6) | n.a. | n.a. | 3/8 (7) | 41 (25) | 34 |
| U2 snRNP (-) | 6/6 (9) | 15 (19) | 26 | 8/8 (8) | 9 (19) | 25 |
| Diacylglycerol kinase (-) | 3/4 (6) | 22 (18) | 27 | 4/4 (5) | 17 (19) | 28 |
| Conserved protein (-) | 0/4 (5) | n.a. | n.a. | n.d. | n.a. | n.a. |
| Dipeptidyl aminopeptid 1 (-) | 0/4 (6) | n.a. | n.a. | n.d. | n.a. | n.a. |
| Hypox-guan phosph transf (-) | 4/4 (8) | 8 (17) | 23 | 4/4 (9) | 1 (15) | 17 |
| Protein kinase (-) | 0/4 (6) | n.a. | n.a. | 0/4 (6) | n.a. | n.a. |
| V-type(+) transl pyrophosph (-) | 1/4 (7) | 21 (13) | 21 | n.d. | n.a. | n.a. |

| | Natural transmission C57Bl/6 | | | 1.000 salivary gland sporozoites i.v. C57Bl/6 | | | 10.000 salivary gland sporozoites i.v. Swiss | | |
|---|---|---|---|---|---|---|---|---|---|
| | mice clearing/ mice infected (prepatency d) | Peak parasitemia cleared [%] (mean d) | Blood stage cleared (mean d) | mice clearing/ mice infected (prepatency d) | Peak parasitemia cleared [%] (mean d) | Blood stage cleared (mean d) | mice clearing/ mice infected (prepatency d) | Peak parasitemia cleared [%] (mean d) | Blood stage cleared (mean d) |
| Dynein heavy chain (-) | n.d. | n.a. | n.a. | 0/4 (4) | n.a. | n.a. | 0/4 (6) | n.a. | n.a. |
| Autophagy-related protein 23 (-) | n.d. | n.a. | n.a. | 0/4 (5) | n.a. | n.a. | 0/4 (5) | n.a. | n.a. |
| Tubulin tyrosine ligase (-) | n.d. | n.a. | n.a. | 0/4 (4) | n.a. | n.a. | 0/4 (6) | n.a. | n.a. |
| Conserved protein (-) | 0/3 (4) | n.a. | n.a. | 0/3 (4) | n.a. | n.a. | n.d. | n.a. | n.a. |
| Protein kinase (-) | 0/6 (5) | n.a. | n.a. | 0/3 (5) | n.a. | n.a. | n.d. | n.a. | n.a. |

**Table S6. Overview of infections with blood stage or salivary gland derived sporozoites from indicated mutants in Swiss and C57Bl/6 mice.**
Natural transmission refers to infection by the bite of 10 infected mosquitoes. Abbreviations: n.a. not applicable, n.d. not determined.

| | 100 iRBC Swiss | | 100 iRBC C57Bl/6 | |
|---|---|---|---|---|
| | dpi reaching peak parasitemia clearing (earliest-mean-latest) | blood stage cleared (earliest-mean-latest d) | dpi reaching peak parasitemia clearing (earliest-mean-latest) | blood stage cleared (earliest-mean-latest d) |
| Plasmepsin IV (-) | 16 | 25 | n.a. | n.a. |
| Autophagy-related protein 23 (-) | 17-20-20 | 22-23-23 | n.a. | n.a. |
| Profilin (-) | n.a. | n.a. | 14-15-16 | 22-23-23 |
| Beta-catenin like protein 1(-) | 17-19-23 | 35-37-39 | n.a. | n.a. |
| Man-1-P-GT (-) | n.a. | n.a. | 21-23-26 | 32-32-33 |
| U2 snRNP (-) | 15-20-22 | 22-26-30 | 16-19-22 | 20-25-30 |
| Diacylglycerol kinase (-) | 16-18-21 | 22-27-31 | 14-19-30 | 24-28-36 |
| Hypox-guan phosph transf (-) | 15-17-19 | 20-23-24 | 14-15-15 | 17 |
| V-type(+) transl pyrophosph (-) | 13 | 21 | n.a. | n.a. |
| Aminopeptidase P (-) | 16-18-20 | 20-21-23 | 14-14-15 | 18-21-23 |
| I-Aminopeptidase (-) | 17-18-19 | 20-21-23 | 14-15-15 | 19-20-20 |

| | Natural transmission C57Bl/6 | | 1.000 salivary gland sporozoites i.v. C57Bl/6 | | 10.000 salivary gland sporozoites i.v. Swiss | |
|---|---|---|---|---|---|---|
| | dpi reaching peak parasitemia clearing (earliest-mean-latest) | blood stage cleared (earliest-mean-latest d) | dpi reaching peak parasitemia clearing (earliest-mean-latest) | blood stage cleared (earliest-mean-latest d) | dpi reaching peak parasitemia clearing (earliest-mean-latest) | blood stage cleared (earliest-mean-latest d) |
| Aminopeptidase P (-) | 12-14-18 | 20-22-24 | 13-17-31 | 17-27-46 | 12-16-19 | 17-23-38 |
| I-Aminopeptidase (-) | 9-15-25 | 17-22-32 | 12-14-18 | 20-22-27 | 15-16-18 | 17-20-23 |

**Table S7. Overview of infections induced either by 100 iRBC or sporozoites with knockout parasite lines used in this study.**

Indicated are the earliest, the mean and the latest day when mice reached peak parasitemia before starting to clear the infection. Also noted is the earliest, mean and latest day when mice became blood stage negative after infection. Abbreviations: n.a. not applicable.

| Immunized through | | | Animals infected/ animals challenged | Average prepatency [d] | Average peak parasitemia [%] | Animals dying/ animals challenged |
|---|---|---|---|---|---|---|
| *Sporozoites i.v.* | *app (-)* | d90 | 6/8 | 10 | 0.9 | 0/8 |
| | | d180 | 3/8 | 10 | 0.5 | 0/8 |
| | | d 360[1] | 4/4 | 8 | 2.3 | 0/4 |
| | *lap (-)* | d90 | 2/8 | 10 | 0.7 | 0/8 |
| | | d180 | 6/8 | 8 | 0.8 | 0/8 |
| | | d 360[1] | 1/4 | 6 | 0.7 | 0/4 |
| *Natural transmission* | *app (-)* | d90 | 2/7 | 12 | 0.4 | 0/7 |
| | | d180 | 2/4 | 10 | 0.3 | 0/4 |
| | *lap (-)* | d90 | 3/8 | 10 | 0.3 | 0/8 |
| | | d180 | 4/4 | 11 | 0.7 | 0/4 |
| *iRBC* | *app (-)* | d90 | 3/4 | 12 | 0.6 | 0/7 |
| | *lap (-)* | d90 | 4/4 | 9 | 0.5 | 0/8 |

[1] mice already challenged at d180 were rechallenged at d360

**Table S8. Summary of all challenged mice and the route of immunization.**

| Gene ID | Gene | Description | RGR | MIS | MFI | # Transf. (gRNAs) | # Transf. Succ. | Growth rate |
|---|---|---|---|---|---|---|---|---|
| PF3D7_ 0725200 | Mago Nashi | mago nashi protein homologue, putative | 0.16 | 1 | -2.2 | 2 (2) | 1 | 0.98 +- 0.09 |
| PF3D7_ 1454400 | APP | Amino-peptidase P | 0.21 | 0.15 | -3.3 | 3 (2) | 0 | - |
| PF3D7_ 1446200 | LAP | M17 leucylamino-peptidase | 0.25 | 0.12 | -3.0 | 2 (2) | 0 | - |
| PF3D7_ 0805700 | FIKK8 | serine/ threonine protein kinase, FIKK family | 0.3 | 1 | -2.7 | 2 (2) | 1 | 0.69 +- 0.02 |
| PF3D7_ 1327600 | NMNAT | Nicotinamide mono Nucleotide adenylyl transferase | 0.41 | 0.75 | -2.9 | 1 (2) | 2 | 1.10 +- 0.11<br>1.42 +- 0.02 |
| PF3D7_ 0828500 | EIF-2B | TIF eIF-2B delta subunit | 0.51 | 0.14 | -2.0 | 1 (2) | 1 | 0.95 +- 0.01 |
| PF3D7_ 1345100 | TRX2 | Thioredoxin 2 | 0.75 | 0.12 | -3.2 | 1 (1) | 1 | 0.8 +- 0.04 |
| PF3D7_ 1148800 | HYP11 | Plasmodium exported protein | no data | 1 | -2.5 | 1 (1) | 1 | 0.94 +- 0.02 |

**Table S9. Growth characterisation of *P. falciparum* knockout lines.**
Relative growth rates of parasite lines. Per mutant, growth rates were normalised to the mean of the respective control line. Welch‘s t-test with correction for multiple comparisons, * $p < 0.05$. Abbreviations: RGR relative growth rate (Zhang 2018), MIS mutagenesis index score (Zhang 2018), MFI mutagenesis fitness index (Zhang 2018), transf. transfection, gRNA guide RNA, succ. Success.
